# Human Naïve Epiblast Cells Possess Unrestricted Lineage Potential

**DOI:** 10.1101/2020.02.04.933812

**Authors:** Ge Guo, Giuliano Giuseppe Stirparo, Stanley Strawbridge, Daniel Spindlow, Jian Yang, James Clarke, Anish Dattani, Ayaka Yanagida, Meng Amy Li, Sam Myers, Buse Nurten Özel, Jennifer Nichols, Austin Smith

**Author notes:** These authors contributed equally (Ge Guo, Giuliano Giuseppe Stirparo). Correspondence (G.G.); (J.N.); (A.S.).

## Abstract

Classical mouse embryology has established a paradigm of early development driven by sequential lineage bifurcations. Accordingly, mouse embryonic stem cells derived from early epiblast have lost the potency to produce extraembryonic trophectoderm. We show in contrast that human naïve epiblast cells readily make trophectoderm. Inhibition of ERK signalling, instrumental in naïve stem cell propagation, unexpectedly potentiates trophectoderm formation, an effect enhanced by Nodal inhibition. Transcriptome analyses authenticate conversion into trophectoderm with subsequent production of syncitiotrophoblast, cytotrophoblast and trophoblast stem cells. Genetic perturbations indicate that NANOG suppresses and TFAP2C enables trophectoderm induction. Consistent with post-implantation progression, trophectoderm potential is extinguished in conventional human pluripotent stem cells, which instead make amnion. Finally, human embryo epiblasts from late blastocysts efficiently generate trophectoderm and differentiated trophoblast. Thus, pluripotent cells in the human embryo retain extraembryonic lineage plasticity and regenerative potential until implantation. Harnessing this unanticipated regulative capacity may be beneficial for assisted reproduction technology.

## INTRODUCTION

Delamination of the epithelial trophectoderm is the first differentiation event in mammalian embryos. Trophectoderm is a cell lineage evolved to mediate blastocyst formation and uterine implantation, and later to produce components of the placenta. Trophectoderm derivatives also provide morphogenetic signals that pattern the early embryo. Following fertilization and early cleavage divisions, blastomeres divide asymmetrically to form trophectoderm and inner cell mass (ICM). Classic studies in mouse embryos have established that the topological segregation of trophectoderm and ICM is rapidly followed by fate restriction, such that by the mid-blastocyst (late 32-cell) stage ICM cells can no longer make trophectoderm (Gardner, 1983; Nichols and Gardner, 1984). Lineage restriction is reflected in the consolidation of distinct molecular identities (Posfai et al., 2017). Subsequently, a second binary fate decision resolves the ICM into epiblast and primitive endoderm (Chazaud et al., 2006; Gardner and Rossant, 1979; Plusa et al., 2008; Saiz et al., 2016). These observations have given rise to a textbook model of sequential lineage bifurcations at the onset of mammalian embryo development (Rossant, 2018).

Mouse embryonic stem (ES) cells are cell lines derived directly from the naïve pre-implantation epiblast (Brook and Gardner, 1997; Evans and Kaufman, 1981; Martin, 1981; Nichols et al., 2009). Over prolonged expansion in vitro they retain global transcriptome proximity to their tissue stage of origin (Boroviak et al., 2014). Functionally, they can contribute massively to all embryo tissues in chimaeras, but do not make appreciable contributions to trophectoderm derivatives (Beddington and Robertson, 1989; Bradley et al., 1984; Nagy et al., 1993; Posfai et al., 2020), in line with the paradigm of early segregation.

Trophectoderm versus ICM determination in the developing mouse blastocyst is underpinned by mutually exclusive and antagonistic expression of transcription factors Oct4 and Cdx2 (Strumpf et al., 2005). ES cells can be made to transdifferentiate into trophectoderm-like cells by either forced expression of *Cdx2* or deletion of *Oct4* (Niwa et al., 2005). Expression of other trophectoderm lineage transcription factors such as Tfap2c (Adachi et al., 2013), or demethylation and up-regulation of Elf5 (Ng et al., 2008), also provoke transdifferentiation into trophectoderm. Detailed characterization, however, has revealed that while cells with morphological features and some markers of trophoblast are obtained, functional phenotypes are not properly established (Cambuli et al., 2014). Moreover, depletion of Nanog, a central component of the ES cell transcription factor network, destabilises naive identity but results in differentiation to primitive endoderm (Chambers et al., 2007; Mitsui et al., 2003), indicating that trophectoderm is not a “default” programme. Thus, trophectoderm lineage restriction appears hard-wired in mouse epiblast and ES cells.

Human pluripotent stem cells (hPSC) (Takahashi et al., 2007; Thomson et al., 1998; Yu et al., 2007) differ from mouse embryonic stem cells and are considered to represent post-implantation epiblast (Rossant, 2015). hPSC have been reported to differentiate into trophoblast-like cells upon treatment with bone morphogenetic protein (BMP) (Amita et al., 2013; Xu et al., 2002). The generation of a pre-implantation lineage by stem cells that have post-implantation identity (Nakamura et al., 2016; O’Leary et al., 2012) is surprising and without developmental precedent. Furthermore, BMP is not involved in trophectoderm specification in the human blastocyst (De Paepe et al., 2019) and the induced cells in vitro do not fulfil stringent criteria for trophoblast identity (Bernardo et al., 2011; Lee et al., 2016). More recently, extended potential hPSCs (hEPSCs) have been described and reported to form trophoblast-like cells, also in a BMP-dependent manner (Gao et al., 2019; Yang et al., 2017). However, the developmental authenticity, either of hEPSCs or of their trophoblast-like progeny, has yet to be ascertained (Posfai et al., 2020).

Culture conditions have now been developed (Guo et al., 2017; Takashima et al., 2014; Theunissen et al., 2014) that support self-renewal of human stem cells with transcriptome proximity to pre-implantation epiblast (Bredenkamp et al., 2019b; Nakamura et al., 2016; Stirparo et al., 2018) and other naïve features (Dong et al., 2019). Their availability provides the opportunity for experimental interrogation of lineage restriction in early human development. Here we investigate naïve cell propensity to produce trophectoderm and show how this differs from the potency of both mouse ES cells and human post-implantation stage PSCs.

## RESULTS

### Human naïve stem cells can enter the trophectoderm lineage

Mouse ES cells self-renew efficiently in the presence of LIF and the MEK inhibitor PD0325901 (PD) (Dunn et al., 2014; Ying et al., 2008). Stable propagation of human naïve stem cells additionally requires the atypical protein kinase C inhibitor Gö6983 and blockade of the Wnt pathway, a culture condition termed PXGL (Bredenkamp et al., 2019b). While investigating the effects of individual inhibitors, we observed that culture PD only resulted in differentiation into flattened epithelial cells (Figure 1A). To determine the character of these differentiated cells we inspected early lineage markers. We did not detect up-regulation of post-implantation epiblast markers that would signify formative transition (Rostovskaya et al., 2019) (Figure 1B). Nor were primitive endoderm factors, *GATA4, PDGFRA* and *SOX17*, expressed (Figure S1A). Instead we observed marked up-regulation of *GATA2* and *GATA3*, transcription factors characteristic of trophectoderm. Strikingly, the other inhibitors in PXGL, XAV939 tankyrase inhibitor and Gö6983, individually reduced, and together completely blocked, expression of *GATA2* and *GATA3* (Figure S1B). We investigated the effect of culture in PD alone on three independent cell lines, including embryo derived and chemically reset naïve cells. Together with up-regulation of GATA2 and GATA3 we saw induction of trophectoderm markers TEAD3 and DAB2 (Figure S1C).

**Figure 1.**
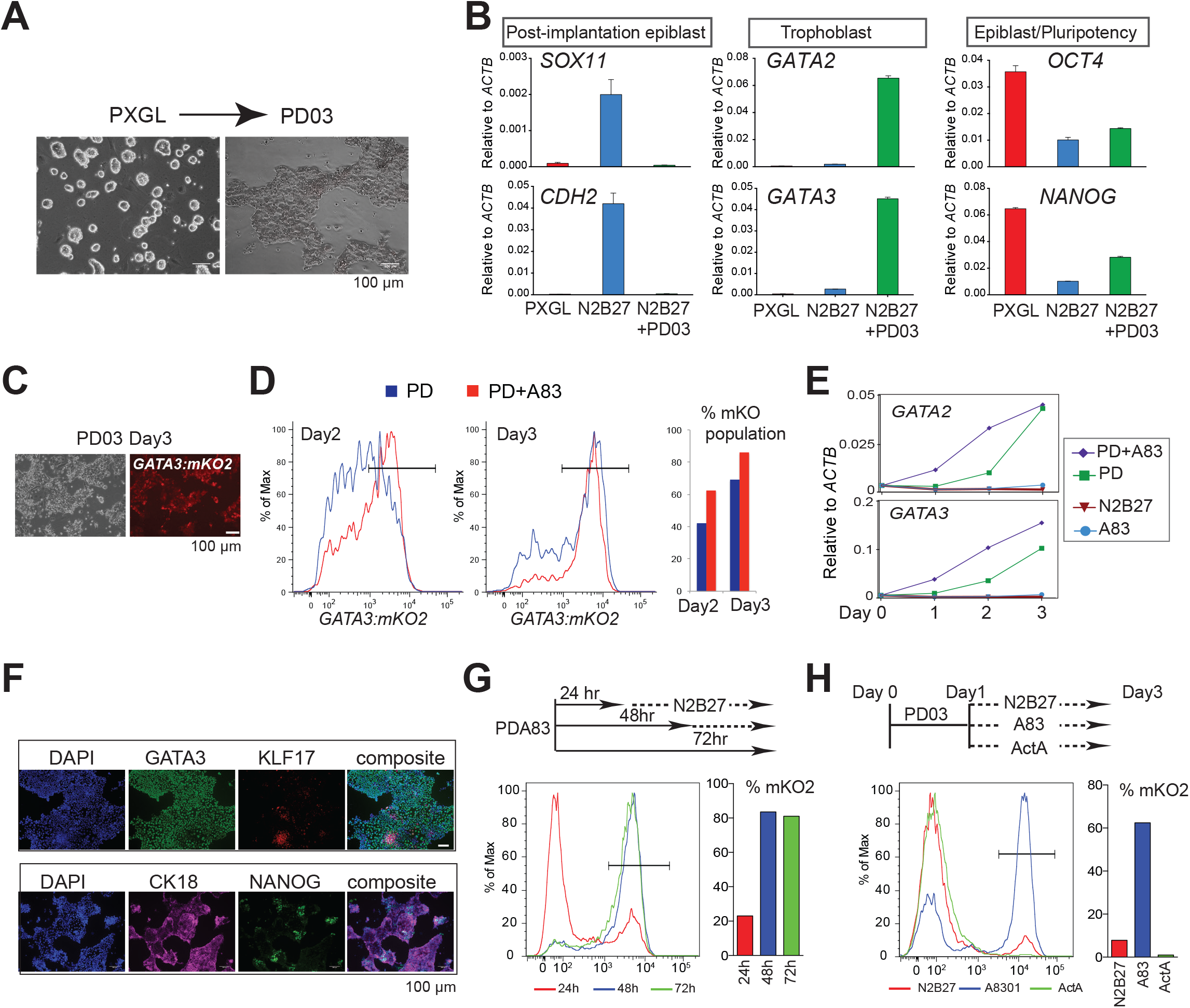
Trophectoderm Formation. A. Images of naïve stem cells and cells differentiating in PD after 3 days. B. RT-qPCR assay for post-implantation epiblast, trophectoderm and core pluripotency markers after 5 days in indicated conditions. Error bars from technical duplicates. C. Phase and fluorescence images of GATA3:mKO2 reporter cells after 3 days in PD only. D. Flow cytometry analysis of GATA3:mKO2 cells exposed to PD+A83 for indicated periods. E. RT-qPCR assay of *GATA2* and *GATA3* expression over time in indicated conditions. Error bars from technical duplicates. F. Immunostaining for trophectoderm markers GATA3 and cytokeratin 18 (CK18) and naïve markers KLF17 and NANOG after 3 days in PD+A83 G. Flow cytometry analysis of GATA3:mKO2 cells treated with PD+A83 for the indicated periods followed by N2B27. H. Flow cytometry analysis of GATA3:mKO2 cells treated with PD for 24h followed by transfer to A83 or Activin A for 48h.

Apparent trophectoderm formation from human naïve stem cells is surprising because mouse ES cells do not generate this lineage without genetic or epigenetic manipulation (Posfai et al., 2020). We cultured mouse ES cells in PD only and did detect weak induction of *GATA3* (Figure S1D). However, neither *GATA2* nor other trophectoderm genes were up-regulated, consistent with inability to enter the lineage. Therefore, the plasticity of human naïve cells in response to PD is species-specific.

To monitor trophectoderm induction we created a *GATA3:mKO2* knock-in reporter line by CRISPR/Cas9 mediated homologous recombination in HNES1 naïve cells (Figure 1C). Fluorescence was barely detected during self-renewal in PXGL or on transfer to N2B27 but was readily apparent in PD only. Naïve stem cells display prominent SMAD2 phosphorylation (Figure S1E) indicative of autocrine Nodal stimulation (Rostovskaya et al., 2019). When Nodal signalling is blocked by the inhibitor A83-01 (A83) (Figure S1F), we noticed that flat differentiated cells appeared at the periphery of colonies. We passaged *GATA3:mKO2* cells in PXGL plus A83. After three passages we detected reporter activation in a significant fraction of cells, coincident with increasing differentiation (Figure S1G,H). We also saw cumulative increases in *GATA2* and *GATA3* mRNAs (Figure S1I). We combined PD and A83 (PD+A83), and saw that mKO2 bright cells appeared earlier and in greater numbers than in PD only, reaching around 80% by day 3 (Figure 1D). We tested A83 alone but observed only rare activation of *GATA3:mKO2* (Figure S1J). mRNAs for *GATA3* and *GATA2* were accordingly up-regulated more rapidly (Figure 1E). Live cell imaging (Figure S1K, Supplemental Movies 1 and 2) showed conversion of HNES1 cells over 60 hours in PD+A83 into an mKO2-positive flat epithelial monolayer of trophectoderm-like cells. Immunostaining after 3 days showed that the majority of cells expressed GATA3 and the epithelial marker cytokeratin 18, exclusive from nests of cells positive for naïve factors NANOG and KLF17 (Figure 1F).

We investigated how long inhibitor treatment is required. We found that 48 hours in PD+A83 was sufficient for robust induction of GATA3:mKO2 and trophectoderm gene expression (Figure 1G). Since A83 alone has little effect, we induced cells with PD for 24 hours then transferred to A83 only. This treatment yielded over 60% mKO2 positive cells (Figure 1H). Conversely, exposure to activin almost entirely suppressed the emergence of positive cells.

These findings indicate that MEK/ERK inhibition is necessary and sufficient to potentiate trophectoderm specification and that NODAL inhibition promotes lineage entry.

### Trophectoderm differentiation and derivation of cytotrophoblast stem cells

During peri-implantation development human trophectoderm gives rise to primary cytotrophoblast cells and syncitiotrophoblast. Cytokeratin 7 (CK7) serves as a pan-trophoblast marker, first expressed weakly in the late blastocyst and subsequently pronounced in cytotrophoblast cells (Deglincerti et al., 2016). During naïve cell differentiation in PD+A83, CK7 was apparent in a few GATA3 positive cells on day 3 then widely and strongly expressed from day 5 (Figure 2A). The syncitiotrophoblast marker β chorionic gonadotrophin (hCGB) was detected in rare clusters of positive cells on day 5 and became more prominent on day 7. RT-qPCR confirmed progressively increasing expression of CK7 together with TEAD3, and presence on day 7 of transcripts for syncitiotrophoblast markers syndecan-1 (SDC1) and chorionic gonadotrophins (Figure 2B)

**Figure 2.**
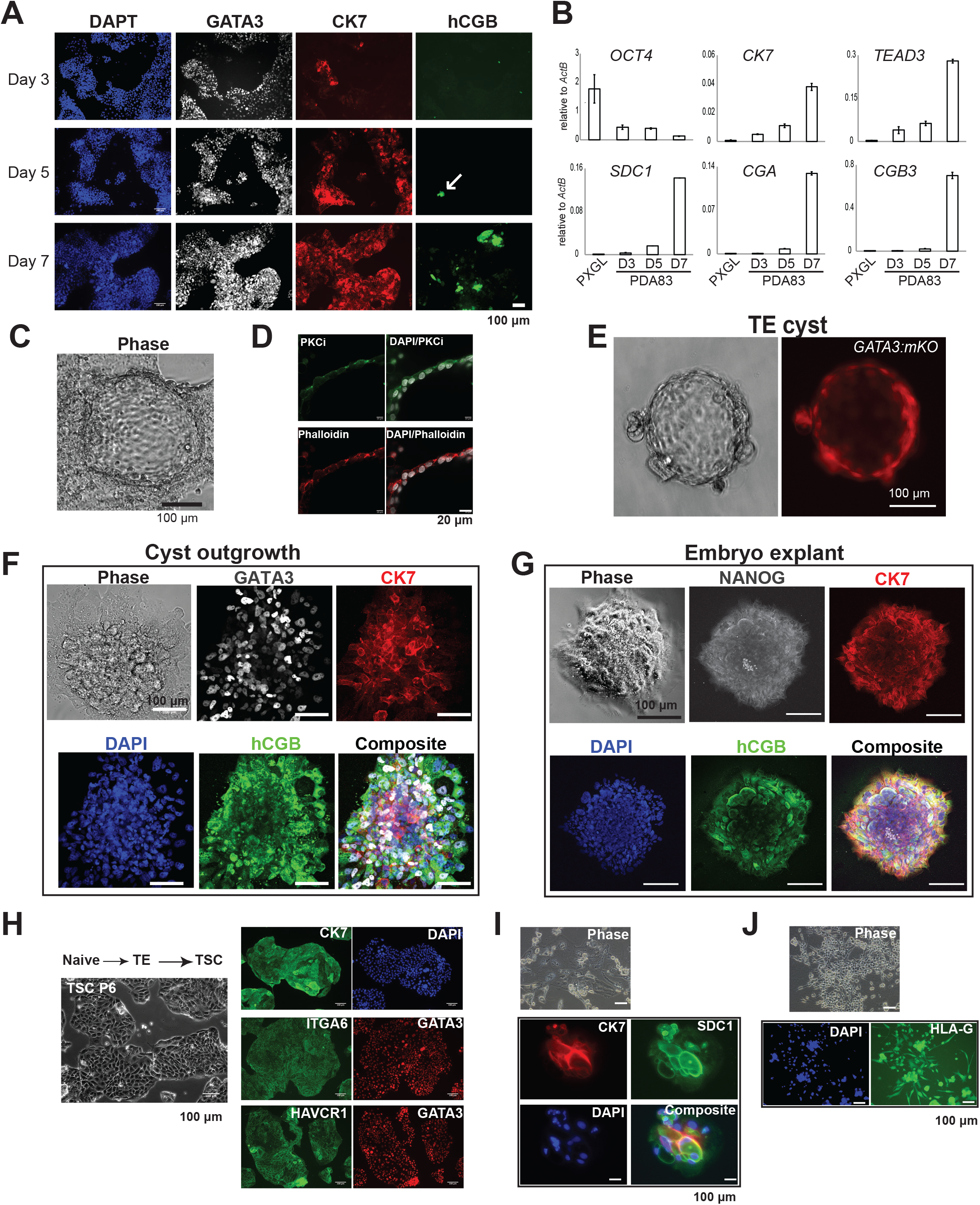
Trophoblast Differentiation and TSC Generation. A. Immunostaining for GATA3, CK7 and hCGB after indicated days in PD+A83 B. RT-qPCR assay of trophoblast marker expression at indicated times. Error bars from technical duplicates. C. Phase image of adherent epithelial cyst formed after PD+A83 treatment for 5 days. D. Confocal image of adherent cyst immunostained for aPKCι. E. Cyst formed in suspension culture in PD+A83 for 4 days. F. Immunostained outgrowth from suspension cyst plated on laminin111-E8 for 5 days in N2B27. G. Immunostained human blastocyst (E6) outgrowth after 5 days in N2B27 on laminin111-E8 H. Phase contrast images of naïve stem cell derived TSCs. I. Immunostaining of naïve stem cell derived TSCs at passage 5. J. Immunostaining of differentiated naïve stem cell-derived TSCs for markers of syncitiotrophoblast (hCGB, SDC1) and extravillous trophoblast (HLA-G).

Trophectoderm is a transport epithelium that mediates formation of the blastocoel cavity by fluid uptake. The emergence in adherent culture of multiple cystic structures indicated functionality of the naïve cell derived trophectoderm (Figure 2C). Consistent with a polarized epithelium, we detected expression of atypical protein kinase C iota (aPCKι) and PAR6B on the apical surface (Figure 2D, S2A). We investigated differentiation in PD+A83 in suspension culture and observed formation of cysts composed of mKO2 positive cells (Figure 2E). When these trophectoderm-like cysts were transferred to dishes coated with laminin111-E8 in N2B27, they attached and formed outgrowths of GATA3 and CK7 positive cells, many of which also expressed hCGB (Figure 2F). The pattern of outgrowth and immunostaining mirrored that in explants of whole blastocysts (Figure 2G). We also detected expression of the extravillous trophoblast marker HLA-G in cyst outgrowths (Figure S2B).

Cytotrophoblast cells from placenta or blastocyst outgrowths can be converted in vitro into human trophoblast stem cells (TSCs) (Okae et al., 2018). We tested whether trophectoderm generated from naïve cells in PD+A83 can give rise to expandable TSCs by replating into TSC medium. Numerous patches of cells with TSC-like morphology emerged within the first passage. After further passage without purification or colony picking, stable and morphologically relatively homogenous epithelial cultures were established, as described for derivations of TSCs (Okae et al., 2018) (Figure 2H). TSCs were derived from different naïve cell lines in two independent experiments and showed similar marker expression as placental TSCs (Okae et al., 2018) (Figure S2B). Immunostaining confirmed expression of CK7, ITGA6, HAVCR1 and GATA3 (Figure 2I). Naïve cell derived TSCs could be induced to differentiate as described (Okae et al., 2018) into hCGB and SDC1 positive syncitiotrophoblast cells and HLA-G expressing extravillous trophoblast (Figure 2J, S2C).

Overall, these observations show that naïve cell derived trophectoderm undergoes progressive differentiation into trophoblast lineage cells in a sequence and pattern that resembles peri-implantation development and that they can readily be converted to TSCs.

### Whole transcriptome analysis of trophectoderm lineage differentiation

We carried out whole transcriptome sequencing over a 5 day time course of naïve cell differentiation in N2B27 alone or with PD, A83 or PD+A83. Duplicate libraries were prepared from both embryo-derived HNES1 and reset cR-H9 cells. Principal component analysis (PCA) aligned samples according to treatment and time along two distinct trajectories (Figure 3A). N2B27 and A83 cultures followed the formative capacitation pathway culminating in the region of density overlay for genes upregulated in conventional hPSCs. PD and PD+A83 samples followed an alternative path extending to the high density area for trophectoderm-enriched genes in the human blastocyst (Petropoulos et al., 2016) (Figure S2A-E). Hierarchical clustering using differentially expressed genes in the embryo substantiated conversion in PD or PD+A83 into a population with trophectoderm features (Figure 3B, S2F). Trophectoderm genes were not upregulated in N2B27 or A83 alone. Figure 3C shows examples of expression dynamics in vitro and in the embryo. Up-regulation of *GATA2, GATA3* and *TBX3* began from day 1, other markers from day 2 or day 3. Some genes (*TFAP2C, TBX3* and *HAVCR1*) prominent in trophectoderm showed appreciable expression in naïve hPSCs (Figure S3G) and were further upregulated in PD and PD+A83.

**Figure 3.**
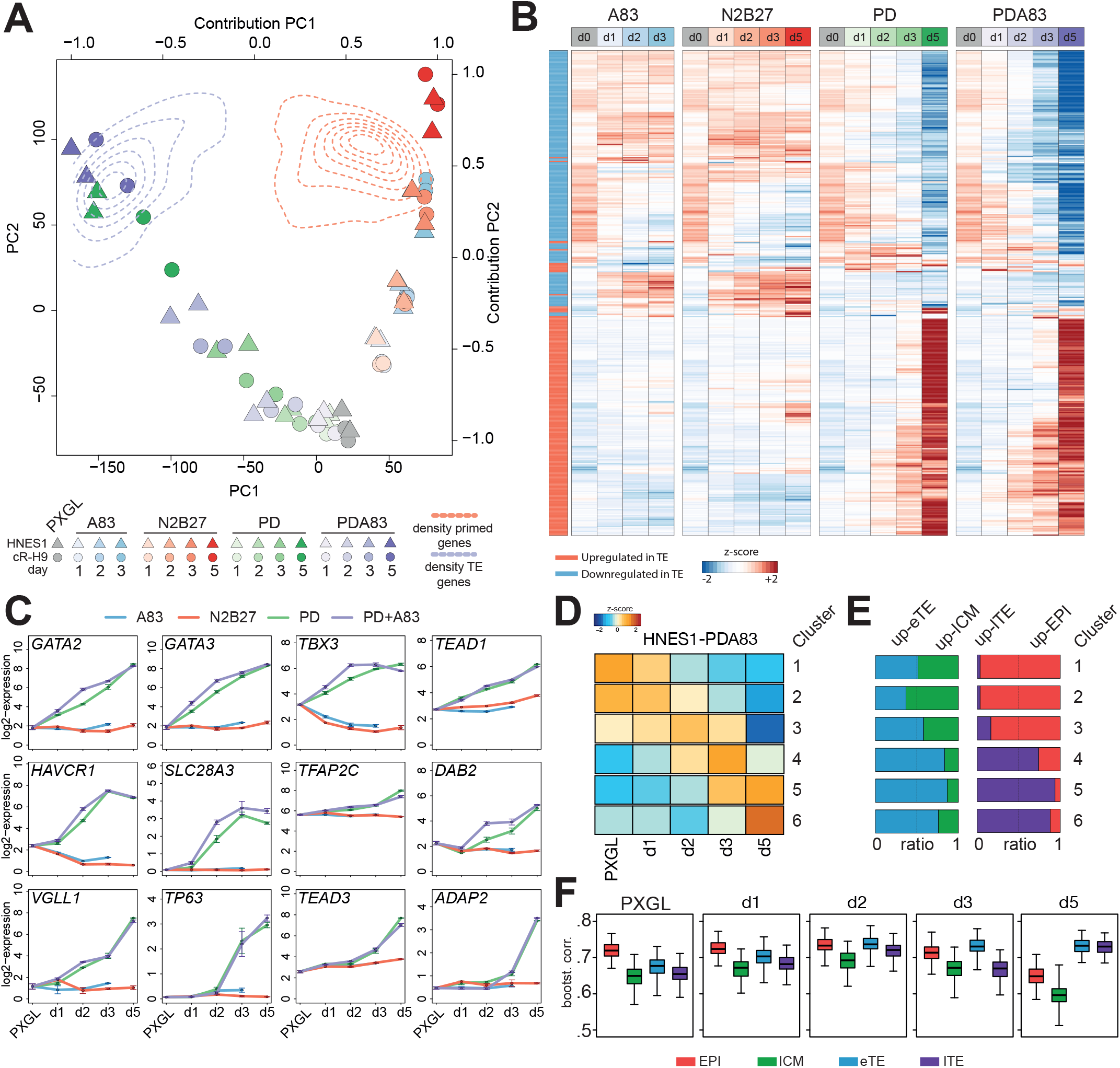
Whole Transcriptome Analysis. A. PCA computed with all expressed protein coding genes (log2 expression >0, n=19450). Two-dimensional kernel density estimation of contribution by genes with enriched expression in trophectoderm (TE), purple dotted lines (late TE vs EPI log2FC > 2, n=409) or in primed hPSCs, red dotted lines (primed VS naïve, Stirparo et al., 2018, log2FC > 2, n=1778). B. One-way hierarchical clustering of differentially expressed genes between late TE (blue) and epiblast (EPI) (red) (top 200 UP and DOWN ranked genes, rank is the product of – log10padj and FC) in HNES1 timecourses in indicated conditions. C. log2 FPKM expression value for selected TE genes in A83, N2B27, PD and PD+A83 conditions. D. Heatmap of zscore centered value for clusters identified in PD+A83 time-course (cluster1: 1787, cluster2: 2594, cluster3: 1872, cluster4: 1227, cluster5: 2055, cluster6: 1839). E. Ratio of modulated genes between eTE/ICM (blue and green) and lTE/EPI (purple and red) in each cluster. F. Bootstrap Spearman correlation (100 iteration, number of genes=50) between PD+A83 time-course (PXGL, d1,d2,d3,d5, log2 expression > 1) and human embryo stages.

For independent assessment comparison with primate embryo development we used transcriptome data from *Macaca fascicularis* (Nakamura et al., 2016). We averaged the scRNA-seq embryo data according to developmental tissue and stage and computed the integrated PCA with orthologous genes. The N2B27 and A83 time course gained similarity to post-implantation epiblast whereas the PD and PD+A83 trajectory related to trophectoderm formation (Figure S3H).

In PD+A83 we identified 6 clusters of dynamically expressed genes (Figure 3D). For each cluster we determined relative representation of profiles of early and late trophectoderm, ICM, and pre-implantation epiblast from the embryo. Clusters 3 to 6 displayed increasing relationship to early trophectoderm compared with ICM and to late trophectoderm compared to epiblast (Figure 3E). We also compared each day of the time course with the embryo samples. Bootstrap Spearman analysis showed an increasing correlation with early trophectoderm from day 1 and with late trophectoderm on day 5. Epiblast correlation declined on day 5 (Figure 3F). Interestingly, correlation with ICM remained low (<0.7) at all time points.

These transcriptome analyses show that human naïve stem cells in PD+A83 do not undergo formative transition but differentiate via a separate path into trophectoderm.

### Single cell transcriptome analysis of differentiation trajectory

To obtain higher resolution of the differentiation trajectory we performed single cell transcriptome analysis using the 10X Genomics platform. We prepared samples on Day 0, 1, 3 and 5 of the PD+A83 time course. A total of 14396 cells passed quality control with >3,000 genes detected. UMAP visualisation showed a relatively continuous and synchronous progression (Figure 4A). Down-regulation of pluripotency markers was reciprocal to up-regulation of trophectoderm genes in the vast majority of cells on day 3 and 5 (Figure 4B,C). A minor fraction of day 5 cells expressed naïve factors and clustered with the day 1 population. Significantly, no cells from day 0 clustered with day 3 or day 5 cells indicating that trophectoderm differentiation does not pre-exist in PXGL culture conditions.

**Figure 4.**
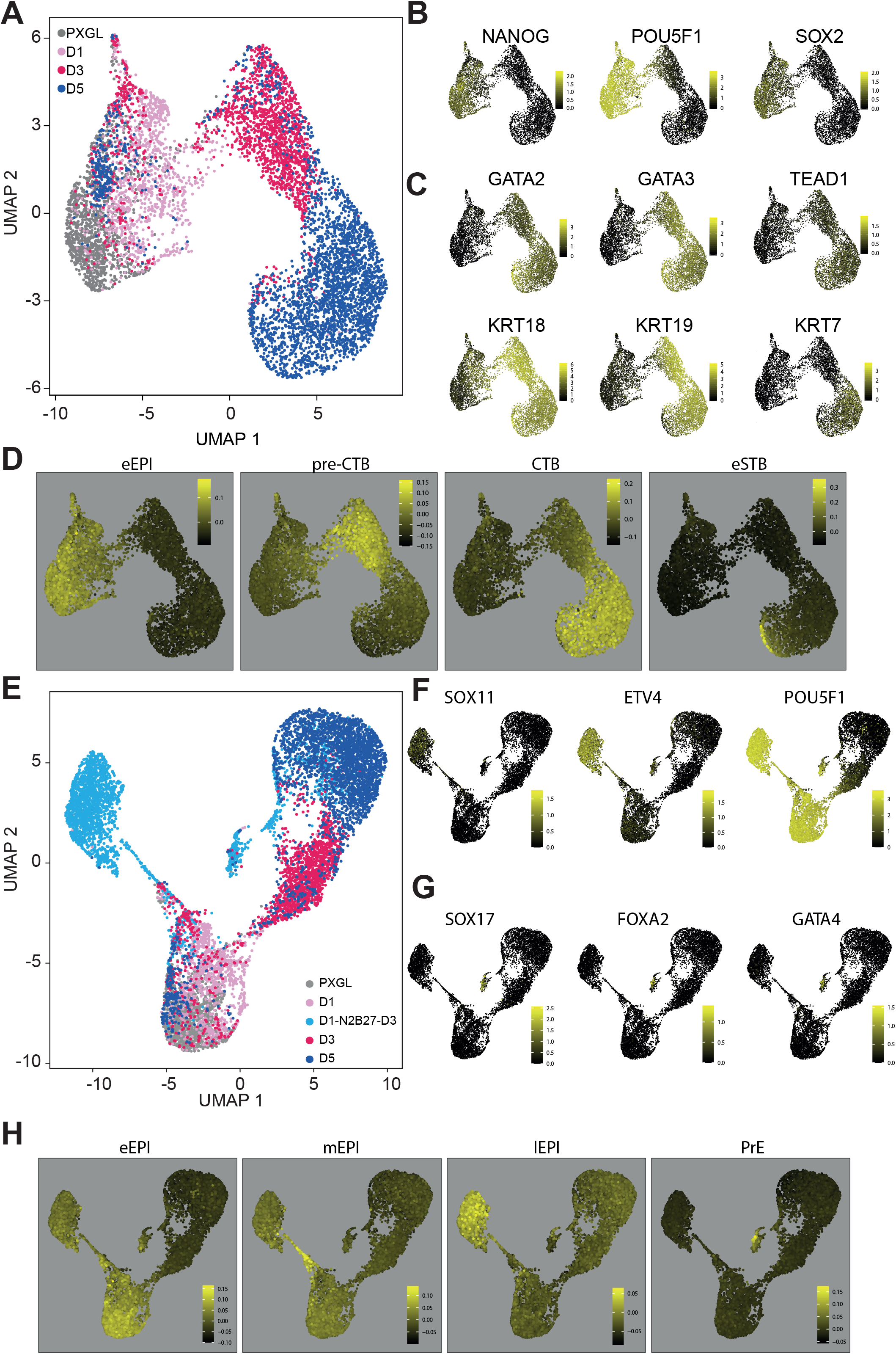
Single Cell Analysis. A. UMAP of PD+A83 time course, colored according to sample day. B. Expression of selected pluripotency markers in A C. Expression of selected trophectoderm and early trophoblast markers in A D. Expression in A of gene lists enriched in indicated human embryo stages (Xiang et al., 2019): eEPI, E6-8 epiblast; preCTB, trophectoderm (E6-7); CTB, cytotrophoblast; eSTB, early syncititiotrophoblast. E. UMAP with addition of cells cultured for 24h in PD+A83 followed by 3 days in N2B27 only. F. Expression of selected post-implantation epiblast markers in E G. Expression of selected primitive endoderm markers in E H. Expression in E of gene lists enriched in indicated human embryo stages (Xiang et al., 2019): eEPI, E6,7,8 epiblast; mEPI, E9,10 epiblast; lEPI, E12,14 epiblast; PrE, primitive endoderm.

We utilized data from extended cultures of human embryos (Xiang et al., 2019) to define gene signatures for tissues and stages, including post-implantation trophoblast types. Computing the distribution of tissue-specific profiles on the UMAP showed conversion from naïve epiblast similarity on day 0 to trophectoderm (termed preCTB by Xiang et al) by day 3 and cytotrophoblast on day 5 (Figure 4D). We also saw that a subset of day 5 cells exhibited features of early syncitiotrophoblast. Inspection of selected trophoblast marker genes substantiated these relationships (Figure S4A), consistent with immunostaining above (Figure 2A)

We also examined the fate of cultures treated with PD+A83 for only 24h then released to N2B27 for 3 days. Integration of this sample (5030 cells) into the UMAP did not affect the major clusters of trophectoderm and cytotrohoblast. However, relatively few of these cells progressed to trophectoderm (Figure 4E). Instead they mainly populated a new cluster that lacked naïve pluripotency factors but expressed general pluripotency and post-implantation epiblast markers (Figure 4F). Interestingly, a stream of cells connecting this cluster with naïve and day 1 cells expressed genes enriched in E8-11 epiblast (Figure S4B), consistent with progression via formative mid-epiblast towards primed late epiblast. A second smaller cluster exhibited a repertoire of primitive endoderm marker genes (Figure 4G, S4C). A few cells from day 3 and day 5 of PD+A83 treatment co-located in this cluster. Distribution of embryo tissue profiles substantiated post-implantation epiblast and primitive endoderm assignations (Figure 4H).

We immunostained cultures treated with PD+A83 for 24 hours then for 2 days with A83 alone or N2B27. In both cases we saw exclusive expression of GATA3, OCT4 or the primitive endoderm marker SOX17 (Figure S4D). Notably, however, the GATA3 population was predominant in A83 whereas the majority of cells were OCT4 positive in N2B27. We surmise that cells treated with PD+A83 for 24h are mostly still flexible in fate choice and can become trophectoderm, primitive endoderm or formative epiblast.

### Genetic perturbation of regulatory transcription factors

In mouse ES cells deletion of *Oct4* or *Sox2* results in up-regulation of *Cdx2* and differentiation to trophoblast-like cells. In contrast, deletion of *Nanog* provokes differentiation to primitive endoderm-like cells with no evidence of trophectoderm (Mitsui et al., 2003). To determine whether these relationships are conserved in human naïve cells we performed targeted mutagenesis using two different CRISPR/Cas9 methodologies.

We first mutated *OCT4, SOX2* and *NANOG* in parental HNES1 cells by transfection with Cas9 and gRNA ribonucleoprotein complexes (RNP). Consistent with their expected essential roles, we observed reduced naïve colony numbers in PXGL for all three genes (Figure S5A). To assess the fate of targeted cells, we performed immunostaining for GATA3 together with the targeted transcription factor on cells either maintained in PXGL or exchanged to N2B27 after 24hrs. On day 5 after transfection, clusters of GATA3 positive cells were apparent in each case (Figure 5A,B). In PXGL depletion of the targeted transcription factor was evident in the cells that were GATA3 positive. RT-qPCR confirmed up-regulation of trophectoderm markers (Figure 5C).

**Figure 5.**
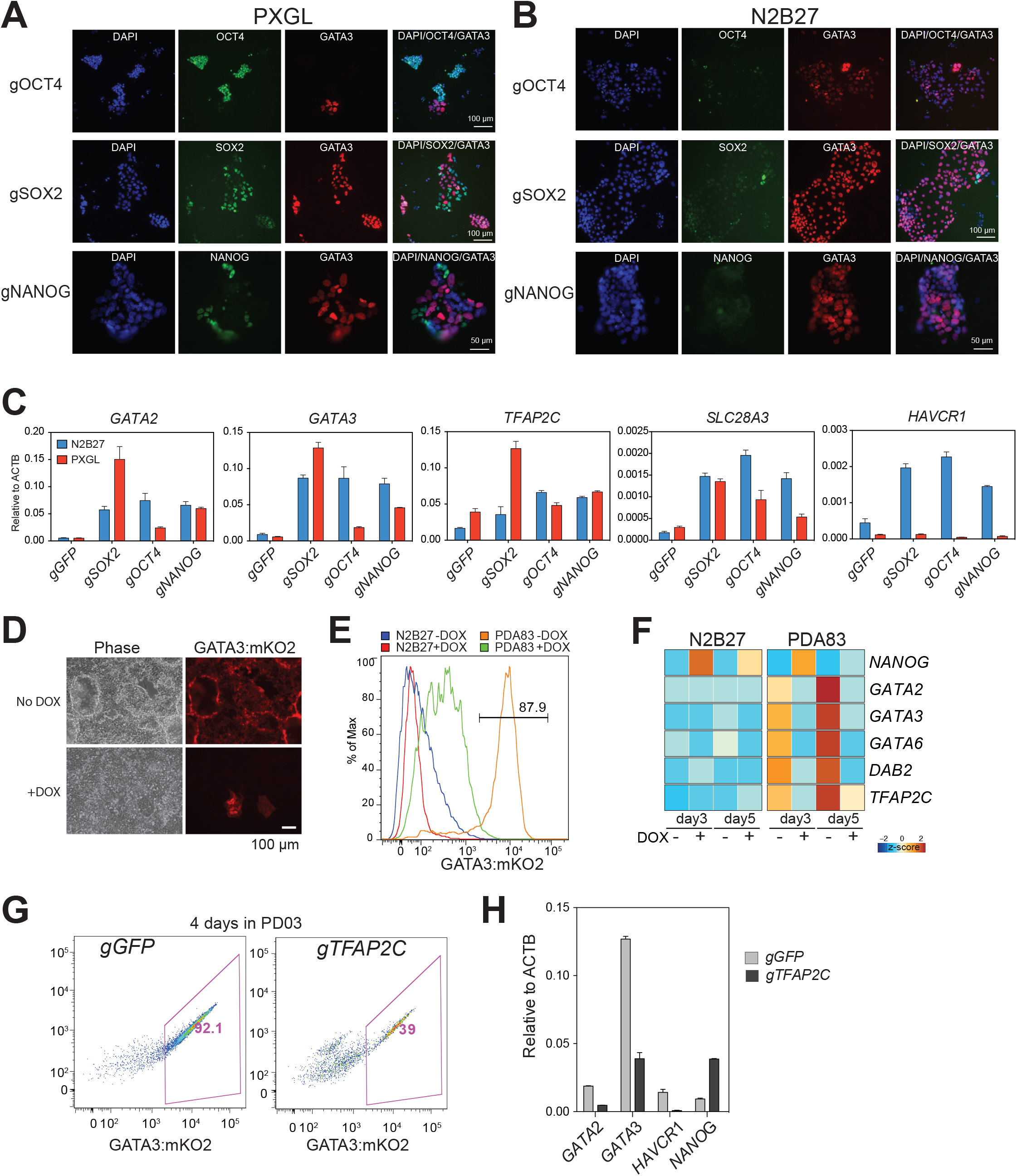
Genetic Perturbations. A. Immunostaining for indicated markers after Cas9/gRNA RNP targeting of *OCT4, SOX2* or *NANOG* in HNES1 cells and maintenance in PXGL for 5 days. B. As A, but culture changed to N2B27 after 1 day. C. RT-qPCR assay of trophectoderm marker expression after Cas9/gRNA RNP targeting of indicated genes in HNES1 cells. Error bars from technical duplicates. D. GATA3:mKO2 cells with or without DOX induction of *NANOG* in PD+A83 for 5 days. E. Flow cytometry histogram of GATA3:mKO2 expression in N2B27 or PD+A83 with or without DOX induction of *NANOG*. F. Heatmap of RT-qPCR gene expression values with and without DOX induction of NANOG in N2B27 or PD+A83. G. GATA3:mKO2 flow cytometry plots after *GFP* (control) or *TFAP2C* targeting by gRNA plasmid transfection and culture for 4 days in PD. H. RT-qPCR assay of marker expression after *GFP* or *TFAP2C* targeting and culture as in H. Error bars from technical duplicates.

RNP transfection efficiency can limit mutation frequency. We therefore introduced a constitutive Cas9 expression construct into the *AAVS1* locus in naïve GATA3:mKO2 reporter cells. We then used piggybac (PB) transposition to integrate gRNA expression constructs containing a selectable marker. After transfection and selection there was a massive reduction in colony numbers in PXGL for all three knockouts (Figure S5B). We saw activation of the mKO2 reporter in N2B27 with or without A83 (Figure S5C). Notably, the *SOX2* knockout had a pronounced phenotype with around 50% and 90% of cells positive for mKO2 in N2B27 and A83 respectively (Figure S5C).

Trophectoderm differentiation in response to *NANOG* targeting is at variance with the phenotype in mouse ES cells, suggesting a function specific to human naïve cells. This prompted us to investigate whether NANOG can suppress trophectoderm formation. We introduced a doxycycline inducible *NANOG* expression vector into GATA3:mKO2 naive cells. Induction of NANOG prevented the appearance of mKO2 positive cells (Figure 5D,E) and suppressed up-regulation of trophectoderm markers in PD+A83 (Figure 5F).

Substantial expression of TFAP2C is a distinctive feature of human naïve stem cells and pre-implantation epiblast cells (Boroviak et al., 2018; Pastor et al., 2018; Stirparo et al., 2018). In mouse, TFAP2C is known as a trophoblast factor (Cao et al., 2015; Choi et al., 2012). It is barely expressed in mouse ICM, early epiblast or ES cells and forced expression provokes trophoblast-like differentiation (Adachi et al., 2013; Kuckenberg et al., 2010). We targeted *TFAP2C* in Cas9 expressing cells. We saw a reduction in colony numbers in PXGL (Figure S5B), consistent with a report that TFAP2C may be required for stable propagation of human naïve cells (Pastor et al., 2018). In contrast to the other transcription factors, however, mKO2 was not elevated. On the contrary, TFAP2C knock-out populations showed reduced production of mKO2 high cells in PD (Figure 5G) and lower expression of trophectoderm markers (Figure 5H). We obtained similar results with the Cas9/gRNA RNP method (Figure S5D).

These observations indicate that OCT4, SOX2 and NANOG suppress trophectoderm while TFAP2C has dual effects, both supporting naïve stem cell self-renewal but also enabling trophectoderm differentiation.

### Differentiation of post-implantation stage hPSCs

There are contested reports that BMP induces conventional hPSCs to form placental trophoblast-like cells (Amita et al., 2013; Bernardo et al., 2011; Lee et al., 2016; Roberts et al., 2014; Xu et al., 2002; Yabe et al., 2016). We investigated whether BMP signalling was required for trophectoderm induction from naïve cells. We found that addition of either BMP or the BMP receptor inhibitor LDN had negligible effect on induction of GATA3:mKO2 in PD+A83 (Figure S6A). Furthermore, BMP signaling is barely detectable in naïve cells compared to primed hPSCs (Figure S6B).

To compare differentiation behaviours in an isogenic setting we converted GATA3:mKO2 cells to conventional hPSC status (Rostovskaya et al., 2019). We then assayed induction of mKO2 in response to PD, PD+A83 or PD+A83+BMP (Figure 6A). For the converted cells, reporter expression was negligible on day 2 in any condition. mKO2 positive cells appeared by Day 3 in PD+A83 or by Day 5 in PD. BMP modestly accelerated these kinetics. Strikingly, reporter levels were at least 10-fold lower relative to differentiation from naïve counterparts. We also noted that A83 or BMP alone induced low expression of mKO2 in a fraction of converted cells but had no effect on naïve cells (Figure S6C). RT-qPCR analysis confirmed log-fold lower up-regulation of GATA3 in conventional compared with naïve hPSCs (Figure S6D).

**Figure 6.**
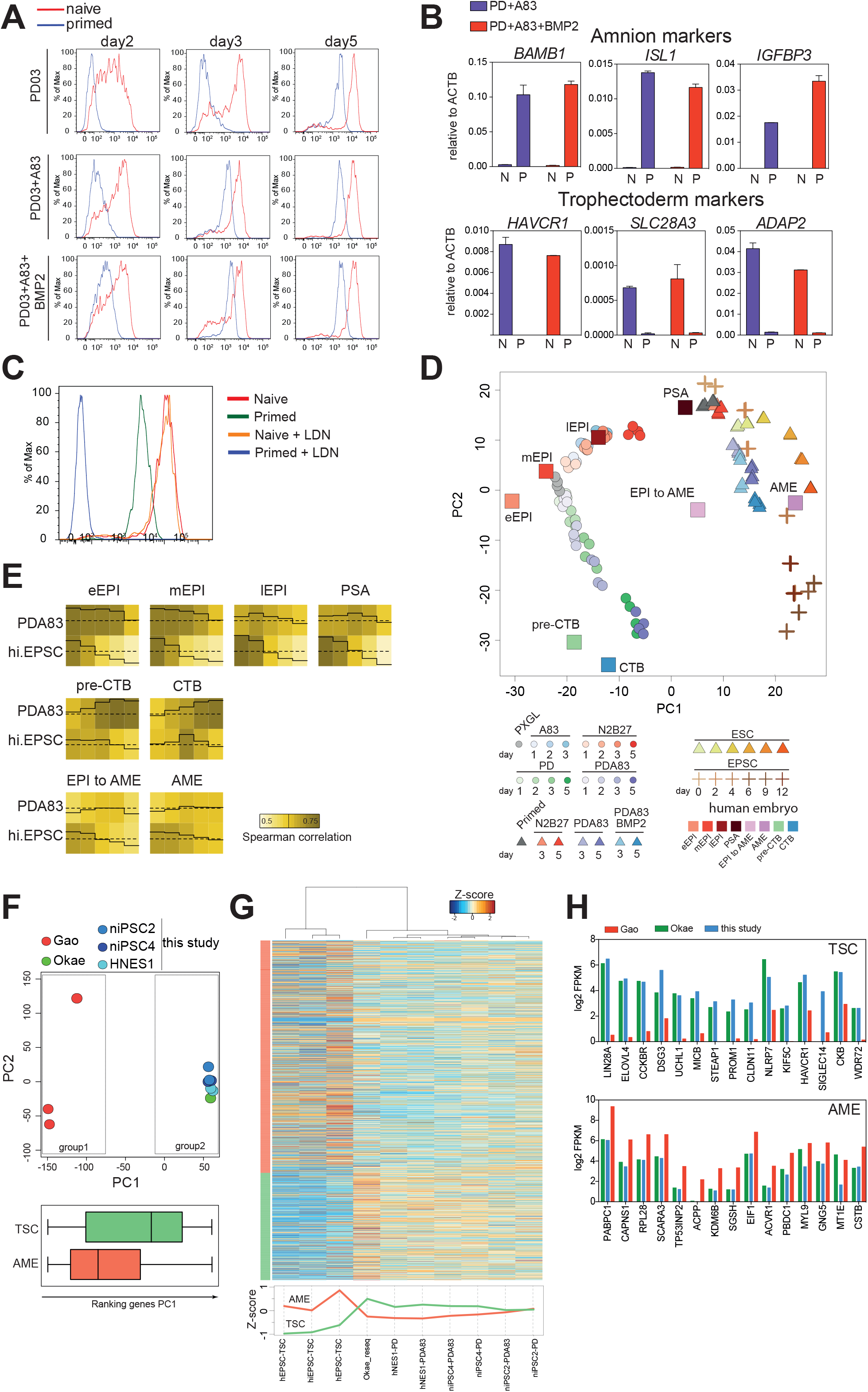
Potency of Naïve Versus Primed hPSCs. A. Flow cytometry analysis of naïve and primed GATA3:mKO2 cells in PD03, PD03+A83 or PD03+A83+BMP2. B. RT-qPCR assay of selected amnion and trophectoderm markers after 5 days culture of naïve (N) or primed (P) cells in PD03+A83 with or without BMP2. Error bars from technical duplicates. C. Flow cytometry analysis of naïve and primed GATA3:mKO2 cells in PD03+A83 with or without BMP inhibitor LDN. D. PCA of A83, N2B27, PD and PD+A83 time courses for naïve cells (HNES1 and cR-H9), average of human embryo stages, hEPSC (d0,d2,d4,d6,d9,d12), primed hPSC (H9, HNES1) time courses in N2B27, PD+A83 and PD+A83+BMP2, and human embryo in vitro development. Computed using 1000 most variable genes between included human embryo stages, with at least log_2_FPKM=1 in one stage). E. Traced heatmap computed with median of bootstrap Spearman correlation (iteration 100, number of genes=50). F. TOP, PCA computed with all expressed genes for hEPSC derivatives (group1), placental TSC and naïve cell derived TSC samples (group2). BOTTOM. Distribution along PC1 of genes enriched for expression in TSC compared with trophoblast lineages in the embryo or in AME compared with other embryo stages (Xiang et al., 2019). G. TOP, Heatmap computed with AME and TSC enriched genes for hEPSC derived TSC-like cells (Gao et al., 2019), placental cytotrophoblast TSCs (Okae et al., 2018) and naïve cell-derived TSCs. BOTTOM, median of Zscore for AME and TSC enriched genes in cell line samples. H. log2 FPKM averaged expression in cells from the indicated studies of top 15 TSC or AME differentially enriched genes.

GATA3 is also expressed in amnion and recently it has been reported that conventional hPSCs differentiate into amnion-like cells in response to BMP (Zheng et al., 2019). In conventional but not naïve cell differentiation we detected upregulation of markers reported by (Zheng et al., 2019) that are also present in early amnion of cultured human embryos (Xiang et al., 2019) (Figure 6B). BMP inhibition with LDN blocked expression of GATA3 and of amnion markers in conventional hPSCs (Figure 6C, S6D) and steered differentiation into the neural lineage (Figure S6E).

More recently, human extended potential stem cells (hEPSC) have been reported to produce trophoblast in response to A83 and enhanced by BMP (Gao et al., 2019). We sought to clarify the relationship between naïve and conventional hPSC or hEPSC lineage trajectories. We examined the transcriptomes of undifferentiated hEPSCs (Gao et al., 2019; Yang et al., 2017) and confirmed that they are distinct from naïve stem cells and similar to conventional hPSCs (Stirparo et al., 2018)(Figure S6F). We compared naïve, conventional and extended potential hPSCs with epiblast stages in the human embryo (Xiang et al., 2019). PCA using differentially expressed genes in the embryo (Figure S6G) corroborated the close relationship between naïve stem cells propagated in PXGL and pre-implantation epiblast (Bredenkamp et al., 2019b; Stirparo et al., 2018), whereas hEPSCs and conventional hPSCs are related to post-implantation epiblast from day 10 onwards.

We compared transcriptomes from hPSC, hEPSC (Gao et al., 2019) and naïve cell differentiation time courses, with human embryo extended culture samples (Xiang et al., 2019), which include post-implantation epiblast to amnion transition and amnion (AME). PCA resolved distinct trajectories for naïve cells compared to conventional hPSCs and hEPSCs (Figure 6D). Naïve cell differentiation in PD+A83 proceeded via similarity to trophectoderm (termed preCTB by Xiang et al.) and culminated in proximity to cytotrophoblast. In contrast, conventional hPSC and hEPSC differentiation was related to early amnion with hEPSCs proceeding further. We extracted genes with enriched expression in amnion-like cells (Zheng et al., 2019) and examined their distribution in the PCA. The intensity overlay was concentrated in the area occupied by the endpoint of hEPSC differentiation (Figure S6H). Analysis using amnion samples from *Macaca* embryo cultures (Ma et al., 2019) produced a similar outcome with the intensity distribution concentrated in the region of differentiated hEPSCs (Figure S6I). We used bootstrap Spearman analysis to examine global correlation between embryo stages and in vitro naïve or hEPSC differentiation. The traced heatmaps show that naïve cell differentiation progressed from high starting correlation with pre-implantation epiblast to similarity with trophectoderm (preCTB) and cytotrophoblast (Figure 6E). In contrast hEPSCs lost high initial relatedness to post-implantation epiblast but did not gain correlation with trophectoderm.

hEPSCs were reported to give rise to TSCs by direct transfer to TSC culture medium (Gao et al., 2019). We compared the transcriptome of hEPSC derivatives (Gao et al., 2019) with placental cytotrophoblast-derived TSCs (Okae et al., 2018) and TSCs derived from naïve cells after PD+A83 induction. PCA computed with all protein coding genes shows naïve stem cell derived and placental TSCs intermingled but well-separated from hEPSC progeny on PC1 (Figure 6F). TSC and AME-enriched genes were differentially distributed along PC1. TSC enriched genes were more highly represented in the naïve stem cell and placenta-derived TSCs whereas hEPSC-derived cells showed higher expression of AME-enriched genes, although many of these were also detected in TSCs (S6J,K). Hierarchical clustering substantiated this finding (Figure 6G). Inspection of the top differentially expressed genes confirmed that TSCs generated from naïve cells expressed TSC markers at comparable levels to placental TSCs and much higher than EPSC derivatives (Figure 6H). Conversely, AME markers were expressed more highly in differentiated EPSCs.

We also examined published transcriptome data for differentiation of conventional hPSCs induced with a combination of BMP, A83 and FGF receptor inhibition (BAP) (Yabe et al., 2016). Bootstrap Spearman correlation analysis showed no significant relationship to naïve cell differentiation in PD+A83 but high correlation with day 4-day 9 of hEPSC differentiation (Figure S6L). We also saw that several amnion enriched genes were expressed in BAP cells similarly to EPSC derivatives (Figure S6M).

Collectively these analyses demonstrate that naive and non-naive hPSCs differentiate along BMP-independent and BMP-dependent trajectories respectively into distinct trophectoderm or amnion-like fates.

### Epiblast in the human blastocyst retains trophectoderm lineage plasticity

Ex vivo culture conditions may corrupt cell identities or alter developmental potential. We therefore examined whether the capacity of naïve stem cells to generate trophectoderm is an authentic feature of epiblast cells in late human blastocysts (E6 and E7) in which primitive endoderm is already specified (Niakan and Eggan, 2013; Roode et al., 2012). Frozen blastocysts (E5 or E6) were thawed and cultured for 24hr for formation of fully expanded blastocysts, some of which commenced hatching (Figure S7A). We confirmed that GATA3 is expressed in all trophectoderm cells and absent from the ICM (Figure S7B). We isolated ICMs for explant culture by immunosurgical removal of trophectoderm (Solter and Knowles, 1975). ICMs were maintained intact and plated on dishes coated with laminin 111-E8 (Kiyozumi et al., 2020) in N2B27 with or without PD+A83. In N2B27 a central mass of relatively undifferentiated cells persisted in most cases but patches of trophectoderm morphology often outgrew. Immunostaining after 5 days showed the expression NANOG in the undifferentiated cell masses and of GATA3, CK7 and hCGB in peripheral cells (Figure 7A). In contrast, in PD+A83 explants invariably differentiated almost entirely into trophectoderm-like cells. GATA3 was expressed throughout the explants together with patches of CK7 and hCGB positive cells by day 5 (Figure 7B). The BMP inhibitor LDN did not impede outgrowth of GATA3 positive cells (Figure S7C, Supplemental Movie 3).

**Figure 7.**
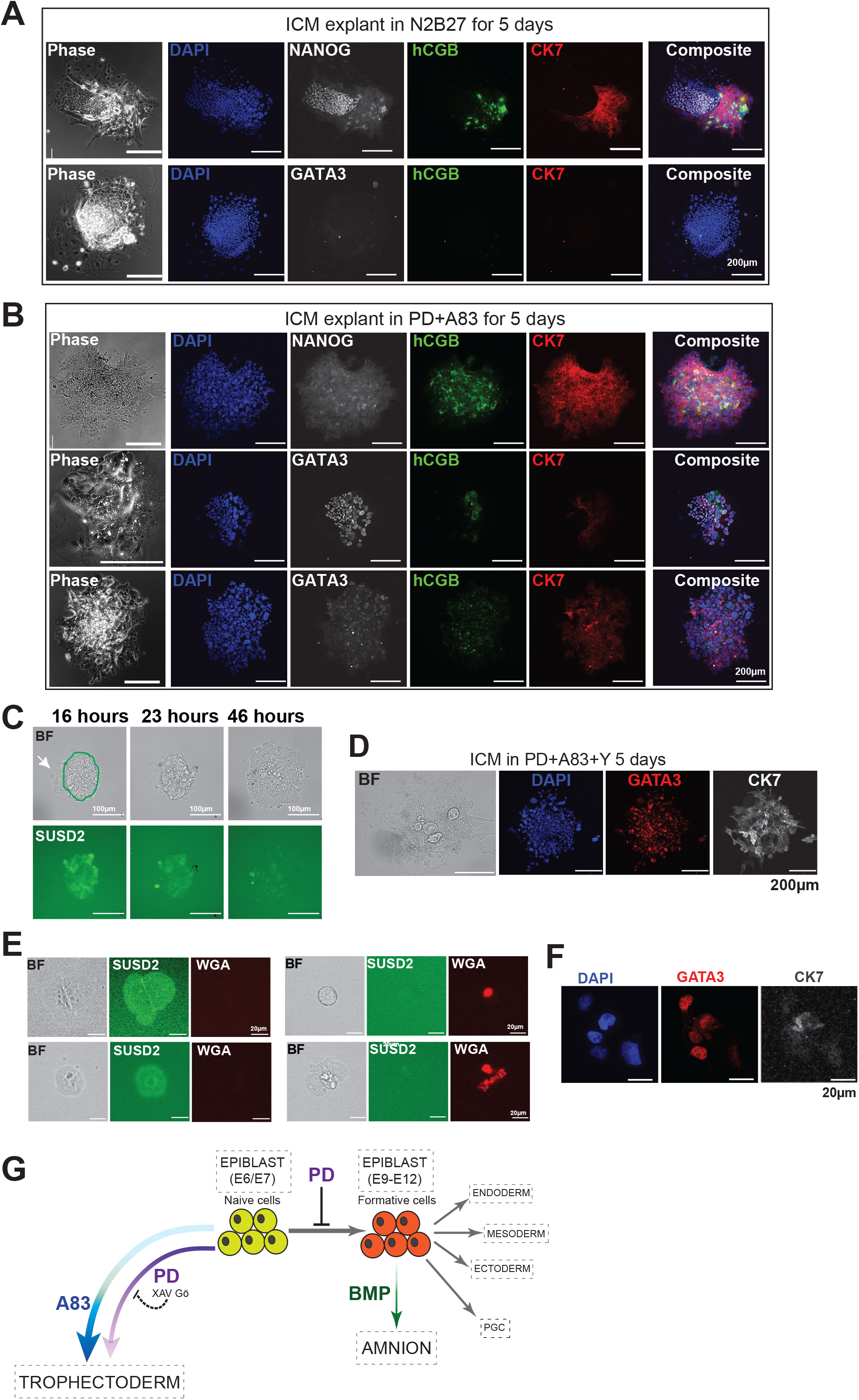
Plasticity of Human Embryo Pre-Implantation Epiblast. A. Phase contrast and immunofluorescence images of ICM explants after 5 days in N2B27 B. Images of ICM explants after 5 days in PD+A83. C. Images of microdissected E6 embryo with epiblast labelled by live cell immunostaining for SUSD2-FITC after culture for 16h. Arrow points to SUSD2 negative cells. Subsequent images show that cells persist as SUSD2 staining fades. D. Immunostaining of microdissected embryo in C after 5 days in PD+A83. E. Images of cells from WGA (Alexa 594) labelled embryos after microdissection, dissociation and plating to attach for 16h before staining for SUSD2 (FITC). F. Immunostained colony formed from cluster of WGA negative, SUSD2 positive cells (upper left panel in E) after culture for 5 days in PD+A83. G. Schematic summary of findings showing that naïve cells can differentiate to trophectoderm or progress to formative pluripotency with switch of lineage competence from trophectoderm to amnion.

These observations suggest that epiblasts from expanded human blastocysts can regenerate trophectoderm. However, the possibility that polar trophectoderm cells may persist after immunosurgery cannot be formally excluded. We therefore adopted an alternative approach. We removed the mural trophectoderm by microdissection and plated the remaining ICM /polar trophectoderm composites. After 16h, we performed live cell immunostaining for the epiblast marker SUSD2 (Bredenkamp et al., 2019a). The antibody labelled ICM cells but not peripheral trophoblast (Figure 7C). Daily imaging showed morphological change with no detachment or appreciable death of ICM cells. After 5 days in PD+A83 the explant comprised almost entirely GATA3 and CK7 positive cells (Figure 7D).

Finally, to confirm epiblast to trophectoderm conversion we incubated intact blastocysts with conjugated wheat germ agglutin (WGA), labelling all outside cells prior to microdissection (De Paepe et al., 2013)(Figure S7D). Dissected WGA-labelled embryo fragments were dissociated using accutase and plated in PD+A83 with addition of Rho-associated kinase inhibitor to safeguard cell viability. Cell positions were registered after attachment at 16h. We performed live cell staining for SUSD2 and captured images, identifying both WGA+SUSD2-negative (trophectoderm) and WGA-SUSD2+ (epiblast) cell clusters (Figure 7E). Cultures were maintained for 5 days in PD+A83 before fixation and immunostaining. Figure 7F shows a tracked colony that arose from a WGA-SUSD2+ cluster. All but one of the cells in this colony was immunopositive for GATA3 (Figure 7G). Faint CK7 staining could also be detected among the GATA3 positive cells.

These findings confirm that human epiblast cells retain plasticity to form trophectoderm. They support a model of pluripotent lineage progression in which trophectoderm potency in human is retained until the formative transition to competence for amnion, germline and germ layer formation (Figure 7H). This sweeping change in developmental capacity is coincident with gain in BMP responsiveness.

## DISCUSSION

Formation of trophectoderm is the first differentiation event in mammalian embryogenesis. In mouse, fate mapping and molecular studies have established a strict lineage bifurcation such that once the blastocyst forms ICM cells become refractory to further trophectoderm specification. Our findings expose a different scenario in human, whereby trophectoderm lineage potential is maintained as the blastocyst matures. Human naïve epiblast stem cells can form trophectoderm with high efficiency and epiblast cells extracted from late human blastocysts robustly regenerate trophectoderm. Lineage restriction is imposed upon formative transition, however. Although post-implantation stage hPSCs can produce epithelial cells superficially similar to trophoblast, global profiling indicates resemblance to amnion, the first lineage to segregate from the embryonic disk in primate embryos (Boroviak and Nichols, 2017). These findings resolve contradictory reports and re-establish a lineage hierarchy consistent with in vivo development. More broadly, our results identify developmental plasticity in the human naïve epiblast that is not precedented by studies in mouse and is likely to confer higher regulative capacity in the human embryo.

Human naïve stem cells are generated and stably maintained in PXGL medium (Bredenkamp et al., 2019b; Guo et al., 2017), which comprises three small molecule inhibitors of MEK/ERK, Wnt, and aPKC signaling respectively. These requirements, which differ from those for mouse ES cell propagation (Dunn et al., 2014; Ying et al., 2008), can now be better understood. In both species, MEK/ERK inhibition prevents formative pluripotency transition to post-implantation epiblast (Smith, 2017). In human, however, PD also promotes differentiation from naïve epiblast to trophectoderm. Wnt and aPKC inhibition block access to trophectoderm, thereby consolidating self-renewal. Thus, triple blockade of signaling input into the gene regulatory network is required to constrain cells in the naïve epiblast state. In addition, autocrine activation of SMAD2 and SMAD3 by Nodal contributes to suppression of trophectoderm. Notably, Nodal and GDF3 are highly expressed in naïve epiblast in the human embryo (Blakeley et al., 2015; Stirparo et al., 2018). In other species, however, exogenous stimulation may facilitate naïve stem cell propagation.

Two recent papers have reported derivation of TSCs from human naïve stem cells by selective amplification of relatively rare cells (Cinkornpumin et al., 2020; Dong et al., 2020). However, it is unclear from those studies how the TSCs arose. In contrast, our findings establish a robust developmental trajectory from naïve epiblast to trophectoderm and thence cytotrophoblast and other trophoblast lineages. We also show that TSCs derived via trophectoderm induction are transcriptomically similar to placental TSCs.

Deletions of core pluripotency factors reveal common and species-specific features. *OCT4* and *SOX2* knockouts trigger trophoblast formation from naïve stem cells in mice and human. *NANOG* deletion also releases trophectoderm differentiation in human naïve cells, whereas mutation in mouse ES cells predisposes to primitive endoderm (Chambers et al., 2007; Mitsui et al., 2003). NANOG may have a conserved function to establish and consolidate naïve epiblast identity, but the outcome of mutation differs due to the lack of trophectoderm restriction in human. Requirement for TFAP2C as a mediator of trophectoderm formation (Cao et al., 2015; Choi et al., 2012) appears conserved. However, in human TFAP2C is also expressed in ICM and epiblast and plays a role in maintenance of naïve stem cell self-renewal (Pastor et al., 2018). We speculate that the dual functionality of TFAP2C may be central to the lineage plasticity of human naïve cells.

There have been several reports that conventional hPSCs and hEPSCs can differentiate into epithelial cells that exhibit some markers of trophectoderm (Amita et al., 2013; Gao et al., 2019; Xu et al., 2002; Yang et al., 2017). However, global transcriptome profiling indicates a trajectory unrelated to pre-implantation development. Instead, differentiation proceeds towards a post-implantation extraembryonic lineage, amnion. Significantly, BMP-signaling is essential for amnion-like differentiation from primed stem cells, whereas trophectoderm induction from naïve cells is insensitive to this pathway. Indeed, naïve cells show negligible SMAD activation when exposed to BMP, indicating that responsiveness to this pathway is only acquired during formative transition.

Our findings demonstrate that human ICM explants from E6 blastocysts have unprecedented plasticity to reform trophectoderm. Mouse ICM loses trophectoderm potency entirely by the mid-blastocyst stage, before epiblast specification (Posfai et al., 2017). By E6 the human ICM has already segregated into primitive endoderm and epiblast (Niakan and Eggan, 2013; Petropoulos et al., 2016; Roode et al., 2012; Stirparo et al., 2018; Xiang et al., 2019). We saw that patches of trophectoderm outgrow from human ICMs in N2B27 alone. This could be attributable to persistence of some polar trophectoderm cells after immunosurgery. However, treatment with PD+A83 caused conversion of almost the entire ICM into trophectoderm and differentiated trophoblast. The ICM origin of regenerated trophectoderm in these conditions was confirmed by prior live cell staining. This finding raises the intriguing possibility that epiblast may contribute continuously to the normal expansion of trophectoderm in the late blastocyst. Alternatively, epiblast plasticity may be reserved for reconstitution after cell loss.

In summary, our results establish that the mouse paradigm of early lineage segregation is not adhered to in human, and that human naïve cells have intrinsic potential for trophectoderm formation. Interestingly, it has been reported that human trophectoderm at E5 can regenerate an ICM population (De Paepe et al., 2013). High regulative flexibility may be an important mechanism for safeguarding human embryos. In the context of assisted conception, this could explain why viable pregnancies can ensue from embryos that are judged morphologically to be of lower quality or incur cell damage during blastocyst freezing and thawing. We speculate that retained trophectoderm potency may be a more widespread feature of early mammalian embryology that has eroded in rodents associated with early implantation and rapid development. Suppressing trophectoderm differentiation may therefore be a common requirement for propagation of naïve pluripotent stem cells.

Recently advances have been made in human embryo culture (Deglincerti et al., 2016; Shahbazi et al., 2016; Xiang et al., 2019). However, availability of human embryos is limited and quality variable. Our study illustrates that human naïve stem cells present a complementary, experimentally convenient, model for delineating the molecular mechanisms of early lineage segregation and uncovering species-specific features. Considerable current interest also focuses on the production of synthetic embryos or blastoids, achieved by combining mouse ES cells together with trophoblast stem cells (Rivron et al., 2018). We suggest that the competency to produce all three primary lineages may enable constitution solely from human naïve stem cells of a blastocyst entity with full developmental potential. With the additional advantage of efficient clonal genome engineering, this would provide an attractive system for elucidating principles of embryo self-organisation.

### Technical Limitations of the Study

The limited numbers of human embryos available for research and their variable quality present major challenges. Our observations are reproducible over multiple experiments but each with only a small number of ICMs. With a more consistent supply of human embryos it would be feasible to implement a live cell labelling approach to investigate whether ICM and epiblast cells contribute continuously to trophectoderm in the unperturbed blastocyst or only in a regenerative context. Furthermore, there is currently only one transcriptome dataset available for extended cultures of human embryos. Future comparison with additional datasets would strengthen our conclusions. In particular, the identity of differentiated cells induced by BMP-based treatments of conventional or extended potential hPSCs should be corroborated as additional transcriptome datasets for primate amnion and other lineages become available.

## Acknowledgments

We are grateful to Kiyotoshi Sekiguchi for laminin-111E8, Hiroaki Okae for hTSCs and Kosuke Yusa and Bahar Mirshekar for the AAVS1-Cas9 targeting vector. We thank Rosalind Drummond and Tao Huang for technical support. Nicholas Bredenkamp, Reo Shoshi and Chonghyun Cha contributed to cell line modification. Vicki Murray and Maike Paramor generated sequencing libraries and Katarzyna Kania and the Cambridge Institute Genomics Facility enabled 1OX sequencing. Peter Humphreys and Darran Clements supported imaging. The Wellcome-MRC Cambridge Stem Cell Institute receives core support from Wellcome and the Medical Research Council (MRC) of the United Kingdom MRC. This research was funded by MRC. AS is a Medical Research Council Professor

## Author Contributions

Conceptualization, GG, AS; Methodology, GG; Investigation, GG, SS, JY, SM, JC, MAL, BNÖ, JN; Formal analysis, DS, GGS; Writing, AS, GG; Supervision JN, GG, AS

## METHODS AND MATERIALS

### Ethics

Use of supernumerary human embryos in this research is approved by the Multi-Centre Research Ethics Committee, approval O4/MRE03/44, and licensed by the Human Embryology & Fertilisation Authority of the United Kingdom, research license R0178.

### Human embryos

Supernumerary frozen blastocysts (E5 or E6) were thawed and cultured in N2B27 medium under mineral oil. The majority of embryos were cultured for 24 hours for development to late blastocysts (E6 or E7). On rare occasions when E5 and E6 embryos had been frozen together, embryos that were already fully expanded on thawing and judged to be E6 were processed immediately. Embryos that failed to expand fully were discarded.

Immunosurgery was performed as described (Guo et al., 2016). In occasional cases when lysis and removal of the trophectoderm could not be assured, embryos were excluded from the study.

For microdissection, embryos were first labelled by incubation with WGA conjugate for 10 minutes and washed in pre-equilibrated N2B27. Mural trophectoderm was excised using a finely drawn Pasteur pipette of internal diameter just larger than the embryo. ICM and polar trophectoderm were dissociated using accutase for 10 minutes, followed by aspiration of individual cells into a drop of N2B27 using a finely drawn Pasteur pipette of diameter just larger than a cell.

### Culture of ICMs and embryo cells

Isolated ICMs were placed intact on laminin-coated plates in N2B27 medium with or without inhibitors. For time-lapse imaging and confocal microscopy, ICMs, ICMs with polar trophectoderm, or dissociated cells were cultured on Ibidi 24-well μ-plates coated with recombinant Laminin-111 E8 (Taniguchi et al., 2009)(kindly provided by Kiyotoshi Sekiguchi). Rho kinase inhibitor Y-27632 was added to dissociated cell cultures. After 16 hours the positions of single cells were registered and live cell SUSD2 immunostaining performed (Bredenkamp et al., 2019a).

### hPSC culture

Naïve and primed hPSCs were cultured in 5% O_2_, 7% CO_2_ in a humidified incubator at 37°C. Cell lines were maintained without antibiotics and confirmed free of mycoplasma contamination by periodic in-house PCR assay.

#### Naïve stem cells

Chemically reset (cR) and embryo-derived (HNES1) naïve stem cells were propagated in N2B27 with PXGL [1μM PD0325901 (P), 2μM XAV939 (X), 2μM Gö6983 (G) and 10ng/mL human LIF (L)] on irradiated MEF feeders as described (Bredenkamp et al., 2019b). Y-27632 and Geltrex (0.5μL per cm^2^ surface area; hESC-Qualified, Thermo Fisher Scientific, A1413302,) were added during replating. Cultures were passaged by dissociation with Accutase (Biolegend, 423201) every 3-5 days.

#### Conventional hPSCs

Conventional primed hPSCs (H9, Shef6) were propagated on Geltrex in Essential 8 (E8) medium made in-house (Chen et al., 2011) or in AFX medium (N2B27 basal medium with 5ng/mL Activin A, 5ng/mL FGF2 and 2μM XAV).

### Differentiation

Human naïve cells were plated in PXGL with Y-27632 on Geltrex or Laminin at a 1:4 to 1:6 ratio. The next day, cultures were washed twice with PBS and medium exchanged to N2B27 with chemical inhibitors or cytokines. Medium was refreshed every day until assaying. Human primed cells were plated in AFX medium on Geltrex at 1:6 to 1:10 ratio and exchanged to assay conditions similarly to naïve cells. Medium was refreshed every day until assaying. Concentrations used in this assay: PD03 1μM, A83-01 1μM, BMP2 50ng/mL, LDN-193189 100nM, Activin A 20ng/mL.

### Capacitation of human naïve cells

Cells were passaged once without feeders in PXGL medium then exchanged into N2B27 containing 2μM XAV for 10 days (Rostovskaya et al., 2019), followed by propagation in AFX medium.

### Generation of GATA3:mKO2 reporter cell line

The *pGG195/GATA3:mKO2* targeting vector was designed to insert an *IresmKO2-FRT-PGKNeobPA* cassette following the stop codon of *GATA3*. 1×10^6^ HNES1 naïve cells were transfected with 3μg *pGG195/GATA3mKO2* and 3ug *px459/GATA3* gRNA using 100 μl Neon transfection kit. G418 (250ng/mL) selection was applied 2 days after transfection for 4 days. Cells were then harvested and transfected with *CAGGS-Flp* plasmid and plated in 2×10cm plates. Clones were picked 7 days after transfection and assayed for mKO2 expression in PXGL and PD03. Genomic DNA was prepared and correctly targeted heterozygous clones were confirmed by PCR amplification of the targeted junction and sequencing.

### Trophoblast stem cell culture

After 3 -5 days treatment with PD+A83, cultures were passaged onto MEF or collagen IV-coated dishes in trophoblast stem cell culture medium (Okae et al., 2018); DMEM/F12 supplemented with 0.1mM 2-mercaptoethanol, 0.2% FBS, 0.3% BSA, 1% ITS-X supplement, 1.5 mg/ml L-ascorbic acid, 50 ng/ml EGF, 2μM CHIR99021, 1.0μM A83-01, 0.8mM VPA and 5μM Y-27632. Cells were passaged by dissociation with TrypLE. Differentiation was induced as described (Okae et al., 2018).

### Inducible NANOG Expression

HNES1/GATA3:mKO2 reporter cells were co-transfected with two *Piggybac* vectors carrying a *Tet3G*-inducible *NANOG* expression cassette and a *CAG-Tet3G-IresZeocin* cassette together with *PBase* plasmid. Two days after transfection, Zeocin (50 ug/mL) was applied for 5 days and individual clones were picked after 7 days. *NANOG* expression was induced with 10-20 ng/mL Doxycycline and assayed by qRT-qPCR.

### CRISPR/Cas9 Knockout

#### Knockout by Cas9/gRNA RNP transfection

TrueGuide synthetic crRNA was purchased from Thermo Fisher Scientific, reconstituted and annealed with tracrRNA in RNA annealing buffer to generate double-stranded RNA duplex. The annealed RNA duplex was diluted in RNA storage buffer to 10μM stock. For each transfection, 1.2μl of the 10μM gRNA duplex was mixed with 300ng Cas9 protein and incubated at room temperature for 15 min before transfection. 10μL Neon transfection kit was used to transfect 1-1.5×10^5^ cells at 1150V, 30ms, 2 pulses. After transfection cells were plated without feeders in PXGL medium with ROCK inhibitor (Y-27632, 10μM). After 24 hours medium was exchanged to N2B27 or other differentiation assay medium for 4 days.

#### Knockout by gRNA plasmid transfection in Cas9 expressing naïve cells

gRNA oligos were synthesized and annealed to double-stranded DNA and cloned behind a U6 promoter (CML32) into a *Piggybac (PB*) vector containing a puromycin resistance gene. gRNA-expression plasmid was transfected together with *PBase* plasmid into HNES1 GATA3:mKO2 cells that had been engineered to constitutively express *Cas9* from the *AAVS1* genomic locus. Following transfection, cells were plated without feeders in PXGL with Y-27632 for 2 days then exchanged to medium for differentiation assay. Puromycin (0.5 μg/mL) was applied for at least 3 days to select cells with stable PB plasmid integration.

### Reverse transcription and real-time PCR

Total RNA was extracted using ReliaPrep kit (Promega, Z6012) and cDNA synthesized with GoScript reverse transcriptase (Promega, A5004) and oligo(dT) adapter primers. TaqMan assays (Thermo Fisher Scientific) and Universal Probe Library (UPL) probes (Roche Molecular Systems) were used to perform gene quantification.

### Immunostaining

Cells were fixed with 4% PFA for 10 min at room temperature and blocked/permeabilised in PBS with 0.1% Triton X-100, 5% Donkey serum for 30 min. Incubation with primary antibodies was overnight at 4°C. Wash was in 0.1% Triton X-100 twice, 10 min each time. Secondary antibodies were added for 1 h at room temperature. Whole embryo and embryo explant staining was performed as described (Guo et al., 2016)

### Microscopy

Wide field images were taken using Leica DMI3000. Confocal images were taken using a Leica SP-2 system. Time-lapse images were taken using a Leica DMI6000 Matrix system fitted with a controlled temperature and CO_2_ chamber. Images were analyzed with Image J software.

### Flow cytometry

Flow cytometry was carried out on CyAn ADP (Beckman Coulter) or BD LSR Fortessa instruments (BD Biosciences) with analysis using FlowJo software. DAPI staining was used to exclude the dead cell population.

### Transcriptome sequencing

For Bulk RNA seq, total RNA was extracted from two biological replicate cultures of each cell line and time point using TRIzol/chloroform (Thermo Fisher Scientific, 15596018), and RNA integrity assessed by Qubit measurement and RNA nanochip Bioanalyzer. Ribosomal RNA was depleted from 1μg of total RNA using Ribozero (Illumina kit). Sequencing libraries were prepared using the TruSeq RNA Sample Prep Kit (RS-122-2001, Illumina). Sequencing was performed on the Novaseq S1 or S2 platform (Illumina).

For 10x Genomics single cell RNA seq, cultures were dissociated with TrypLE™ Express Enzyme at 37 degree for 10 min. Single cell populations were sorted using a flow cytometer based on forward/side scatter into PBS with 0.04% BSA. Single cell libraries were captured using Chromium Single Cell 3’ Reagent Kits and sequenced on a Novaseq 6000 sequencer. Approximately 3000-5000 cells were captured for each time point.

## QUANTIFICATION AND STATISTICAL ANALYSIS

Alignment was performed using the Genome build GRCh38 and STAR (Dobin et al., 2013) were used for aligning reads. Ensembl release 96 was used to guide gene annotation. After removal of inadequate samples, we quantified alignments to gene loci with htseq-count (Anders et al., 2014) based on annotation from Ensembl 96. Principal component, differential expression and cluster analyses were performed based on log2 expression values computed with custom scripts, in addition to the Bioconductor packages DESeq (Anders and Huber, 2010), FactoMineR (Lê et al., 2008) and MFuzz (Kumar and M, 2007). Gene density that contributed to PCA plots were calculated using kernel density estimation (MASS R package (Venables and Ripley, 2002)

For global analyses, we considered only genes with log2 expression > 0 (unless otherwise indicated) in at least one condition, not expressed genes were always omitted. Euclidean distance and average agglomeration methods were used for cluster analyses unless otherwise indicated. Human transcription factor and co-factors were downloaded from (http://bioinfo.life.hust.edu.cn/AnimalTFDB/). Time courses for H1.ESC, H1.EPSC and hiEPSC (Gao et al., 2019) were downloaded from array express (E-MTAB-7253) and re-aligned.

For 10x analyses Cellranger-3.1.0 count (Zheng et al., 2017) was run using default parameters and Cellranger’s prebuilt human reference genome ‘refdata-cellranger-GRCh38-3.0.0’, producing counts matrices which were merged and analysed using Seurat 3.1.5 (Stuart et al., 2019) in R. A bimodal distribution of the number of genes expressed per cell was observed therefore all cells expressing fewer than 3000 genes or more than 10% mitochondrial genes were removed. The remaining cells were log-normalized via the division of each cell’s feature counts by the cell’s total counts, which were then multiplied by a scale factor of 10,000 and finally natural-log (log1p) transformed. The top 2000 most variably expressed genes within the dataset were identified, scaled, and centered for use in initial dimensionality reduction with PCA prior to further non-linear dimensionality reduction using UMAP. Cells visualised in UMAP plots were colored according to individual marker gene expression values and similarity to cell type-specific gene expression signatures, scored using Seurat’s AddModuleScore function. Z-scores were used to plot heatmaps and dotplots of marker expression.

The Cynomolgus monkey dataset (Nakamura et al., 2016) was kindly provided by Dr. Nakamura as an RPM table.

Expression values of ESC cultured in different human naïve/primed conditions and pre-implantation embryo single cell RNAseq were downloaded from (Stirparo et al., 2018). Embryo cells expressing less than 6000 genes with log2 expression > 1.5 were excluded. EPI, PrE and TE markers for cell stratification were selected according to Stirparo et al., 2018.

Human in vitro pre-gastrulating single cells were downloaded from Xiang et al., 2019, GEO accession number GSE136447. High variable genes across EPI cells were computed according to the methods described (Boroviak et al., 2018; Stirparo et al., 2018). A non-linear regression curve was fitted between average log2 FPKM and the square of coefficient of variation (log CV^2^); then, specific thresholds were applied along the x-axis (average log2 expression) and y-axis (log CV^2^) to identify the most variable genes. Cells expressing less than 6000 genes with log2 expression >1.5 were excluded from analysis.

Human amnion-like genes were downloaded from (Zheng et al., 2019) (Supplement Table 1, AMLC genes) and we selected only genes >1 expression in amnion. Early and late AMLC genes in cynomolgus monkey were downloaded from (Ma et al., 2019)(Supplement Table 6).

## DATA AND CODE AVAILABILITY

RNA-seq data from this study are being deposited in Gene Expression Omnibus.

