## Supplemental figures and legends for "Human Naïve Epiblast Cells Possess Unrestricted Lineage Potential"

#### SUPPLEMENTAL FIGURE LEGENDS

**Figure S1. Trophectoderm formation, Relates to Figure 1**

A. RT-qPCR assay for primitive endoderm markers after 5 days in indicated conditions.
B. RT-qPCR assay for naïve and trophectoderm markers after 5 days in indicated conditions.
C. RT-qPCR assay for trophectoderm markers after 5 days in N2B27 or PD03 of human naïve PSC lines; HNES1, cR-NCRM2 and cR-Shef6
D. RT-qPCR assay for core pluripotency and trophectoderm marker expression in mouse ES cells cultured for 3 days in indicated conditions.
E. Phospho-Smad2/3 immunoblot for conventional H9 hPSCs and naïve cells in indicated conditions.
F. Phospho-Smad2/3 immunoblot for naïve cells in absence or presence of A83
G. Phase and fluorescence images of GATA3:mKO2 cells in PXGL with or without A83 for 4 passages. Scale bar, 100μM
H. Flow cytometry analysis of GATA3:mKO2 cells in PXGL with or without A83 for 4 passages.
I. RT-qPCR assay for *GATA2* and *GATA3* expression in human naïve cells cultured in PXGL with A83. X-axis indicates passages in PXGL with A83.
J. Flow cytometry analysis of GATA3:mKO2 cells in N2B27 or with A83 for 3 days.
K. Phase contrast and fluorescence time lapse stills of GATA3:mKO2 cells in PD+A83. (See also Supplemental Movies 1 and 2).

**Figure S2. Trophectoderm differentiation and cytotrophoblast stem cells, Relates to Figure 2**

A. Wide field images of cysts formed in adherent culture in PD+A83 immunostained for aPKCι and PAR6B.
B. Two examples of suspension cysts outgrown for 3 days in N2B27 and immunostained for CK7 and extravillous trophoblast marker HLA-G.
C. RT-qPCR analysis of gene expression in placental and naïve stem cell derived TSCs. Naive stem cell derived TSCs include two independent cultures derived from HNES1 and one each from naïve iPSC lines niPSC2 and niPSC4. CT27 is a placental cytrophoblast TSC line (Okae et al, 2018). Error bars from technical duplicates.
D. RT-qPCR analysis of gene expression during differentiation of placenta and naïve stem cell derived TSCs. Error bars from technical duplicates.

**Figure S3. Whole transcriptome analysis, relates to Figure 3**

A. One-way hierarchal clustering of early blastocyst (E5) single cells (Petropoulos et al., 2016) computed with lineage marker genes (Stirparo et al., 2018).
B. One-way hierarchal clustering of E6 and E7 single cells (Petropoulos et al., 2016) computed with lineage marker genes (Stirparo et al., 2018).
C. PCA computed for all the genes expressed in early ICM (cluster 4, FIG.S2A), early TE (cluster 5, Figure S2A), epiblast (Stirparo et al., 2018) and late TE (cluster 2, Figure S2 B, C, D). In red, cells expressing more than 6000 genes.
D. PCA computed as in C for filtered cells expressing >6000 genes (log2expression >0, n=18694)
E. PCA for filtered cells, computed with differentially expressed genes in human embryo (n= 4507).
F. As Figure 3B for cR-H9 cells.

G. PCA computed with all orthologues (average macaque dataset, ICM, EPI, postE, postL, early TE, late TE and post PA.TE, Nakamura et al., 2016) of expressed protein coding genes (log2 expression in time-course > 0 & orthologues, n= 12992).

**Figure S4. Single cell analysis, relates to figure 4**

A. Expression of cytotrophoblast, syncitiotrophoblast and extravillous trophoblast lineage markers on Figure 4A UMAP
B. Expression of mid (E9-10) and late (E12-14) post-implantation epiblast markers on Figure 4A UMAP
C. Expression of additional primitive endoderm markers on Figure 4E UMAP
D. Immunostaining of cells cultured in PD+A83 for 24h followed by 48h in A83 or N2B27.

**Figure S5. Genetic perturbations, relates to Figure 5**

A. Alkaline phosphatase (AP) staining after indicated Cas9/gRNA RNP transfection and culture in PXGL on MEF for 4 days. Separate experiments were performed on parental HNES1 or HNES1-GATA3:mKO2 cells. Controls were transfected with GFP gRNA.
B. AP staining after indicated gRNA plasmid transfection in Cas9 expressing HNES1-GATA3:mKO2 cells. Cells were maintained in PXGL on MEF with puromycin selection for 7 days. Controls were transfected with GFP gDNA.
C. Flow cytometry analysis of GATA3:mKO2 cells after gRNA plasmid transfection and culture for 4 days in N2B27 alone or with A83.
D. Flow cytometry analysis of GATA3:mKO2 expression after Cas9 RNP transfection with GFP or TFAP2C gRNA.

**Figure S6. Fates of naïve versus primed stem cells, relates to Figure 6**

A. Flow cytometry analysis of naïve GATA3:mKO2 cells in indicated culture conditions for three days. LDN, BMP receptor inhibitor LDN-193189.
B. Phospho-Smad1/5 immunoblot on naïve cells and primed HNES1 cells in indicated conditions. P, PD03; B, BMP2, number indicates BMP2 concentration, ng/ml; L, human LIF.
C. Flow cytometry analysis of naïve and primed GATA3:mKO2 cells in indicated culture conditions for 5 days.
D. RT-qPCR assay of trophectoderm and amnion markers in human naïve and primed cell differentiated in PD+A83 with or without LDN for 5 days. Error bars from technical duplicates.
E. RT-qPCR assay for neural markers in naïve or primed PSCs differentiated in indicated conditions for 5 days with or without LDN. Error bars from technical duplicates.
F. PCA of human naïve and conventional primed cells (Stirparo et al., 2018), naïve cells from this study, and hEPSCs (Gao et al., 2019; Yang et al., 2017) computed using most variable genes, log2expression > 1, cv>0.5, n=3510.
G. 3D PCA of human naïve and conventional primed cells, naïve cells from this study, hEPSCs, and human embryo in vitro development (Xiang et al., 2019), computed with variable genes in embryo development, n=1517.
H. Two-dimensional kernel density estimation of amnion genes from human differentiation in vitro (expression >1, Zheng et al., 2019) in Figure 6D PCA.
I. Two-dimensional kernel density estimation of amnion genes from *macaca* embryo cultures (Ma et al., 2019) in Figure 6D PCA.

J. Density of placental cytotrophoblast TSC (Okae, 2019)(left) and AME (right) enriched genes in Figure 6F PCA.
K. Violin plot of Zscore for group 1 and group 2 samples for TSC and AME enriched genes.
L. Bootstrap Spearman correlation (iteration 100, number of genes=50) of naïve or EPSC differentiation timecourses with differentiated populations induced by BAP treatment.
M. As Figure 6H with addition of BAP cell expression values.

**Figure S7. Late blastocyst ICM explants, relates to Figure 7**

A. Phase images of human blastocysts cultured for 24h and classified as fully expanded (E6 or E7).
B. Immunostaining of E6 human blastocyst for GATA3. White dashed line demarcates ICM population.
C. Time lapse stills at beginning and end of 48h ICM outgrowth culture in PD+A83 plus LDN with immunostaining for GATA3 and SOX2 at endpoint.
D. WGA (Alexa 594 conjugate) labelling of intact E6 blastocysts. Scale bar, 200μM.

**Supplemental Movies 1 & 2**

Phase contrast and fluorescence time lapse of GATA3:mKO2 cells cultured in PD+A83, 60h.

**Supplemental Movie 3**

Time lapse of ICM explant in PD+A83 with LDN, 48hrs

### Figure S1

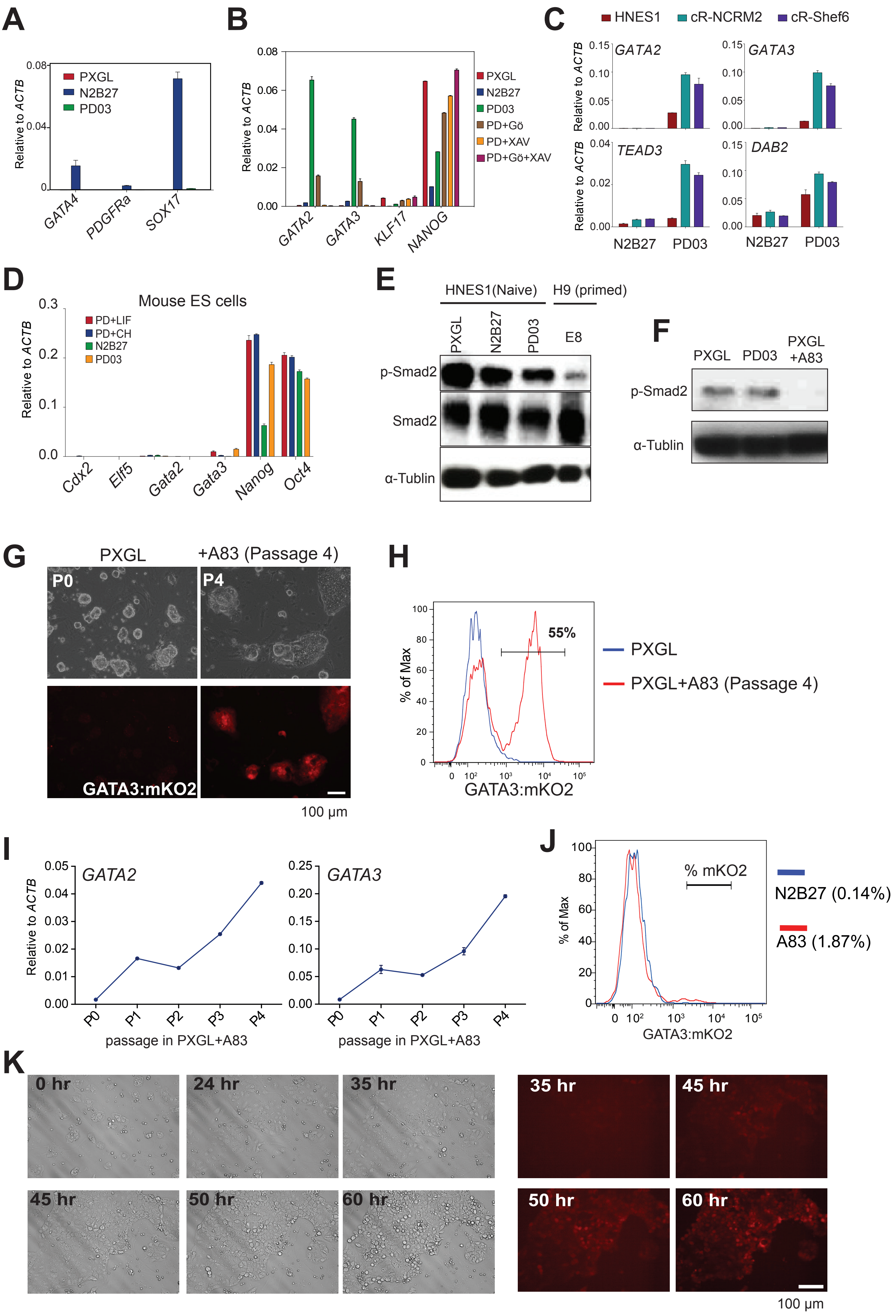

A
Relative to ACTB
PXGL
N2B27
PD03
GATA4
PDGFRa
SOX17
B
Relative to ACTB
PXGL
N2B27
PD03
PD+Gö
PD+XAV
PD+Gö+XAV
GATA2
GATA3
KLF17
NANOG
C
HNES1
cR-NCRM2
cR-Shef6
GATA2
GATA3
TEAD3
DAB2
Relative to ACTB
N2B27
PD03
D
Mouse ES cells
Relative to ACTB
PD+LIF
PD+CH
N2B27
PD03
Cdx2
Elf5
Gata2
Gata3
Nanog
Oct4
E
HNES1(Naive)
H9 (primed)
PXGL
N2B27
PD03
E8
p-Smad2
Smad2
α-Tublin
F
PXGL
PD03
PXGL +A83
p-Smad2
α-Tublin
G
PXGL
+A83 (Passage 4)
P0
P4
GATA3:mKO2
100 μm
H
% of Max
55%
PXGL
PXGL+A83 (Passage 4)
GATA3:mKO2
I
GATA2
GATA3
Relative to ACTB
passage in PXGL+A83
J
% of Max
% mKO2
N2B27 (0.14%)
A83 (1.87%)
GATA3:mKO2
K
0 hr
24 hr
35 hr
45 hr
50 hr
60 hr
100 μm

Figure S2

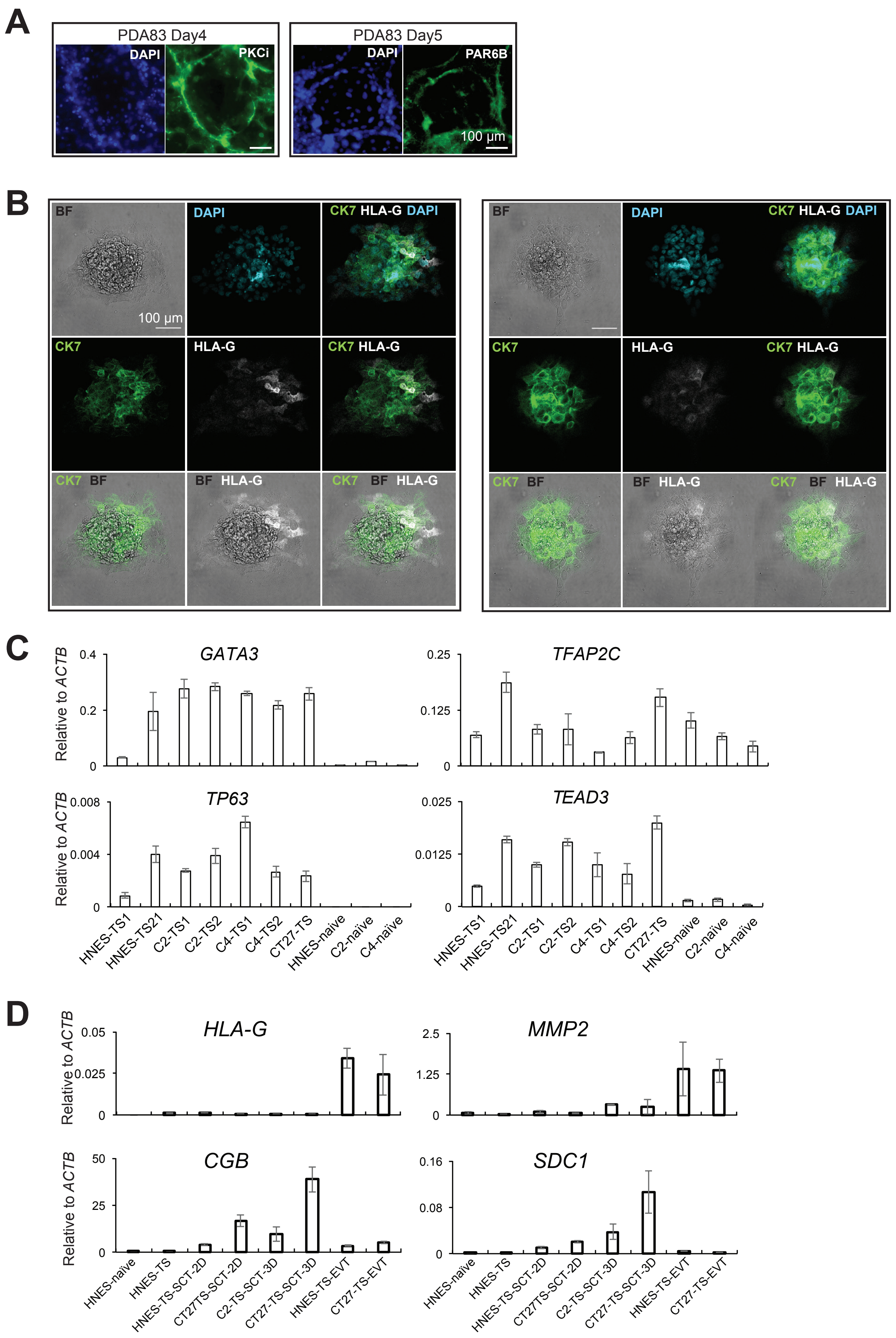
A
PDA83 Day4
DAPI
PKCi
PDA83 Day5
DAPI
PAR6B
100 µm
B
BF
DAPI
CK7 HLA-G DAPI
100 µm
CK7
HLA-G
CK7 HLA-G
CK7 BF
BF HLA-G
CK7 BF HLA-G
C
GATA3
TFAP2C
TP63
TEAD3
Relative to ACTB
HNES-TS1
HNES-TS21
C2-TS1
C2-TS2
C4-TS1
C4-TS2
CT27-TS
HNES-naïve
C2-naïve
C4-naïve
D
HLA-G
MMP2
CGB
SDC1
HNES-naïve
HNES-TS
HNES-TS-SCT-2D
CT27TS-SCT-2D
C2-TS-SCT-3D
CT27-TS-SCT-3D
HNES-TS-EVT
CT27-TS-EVT

Figure S3

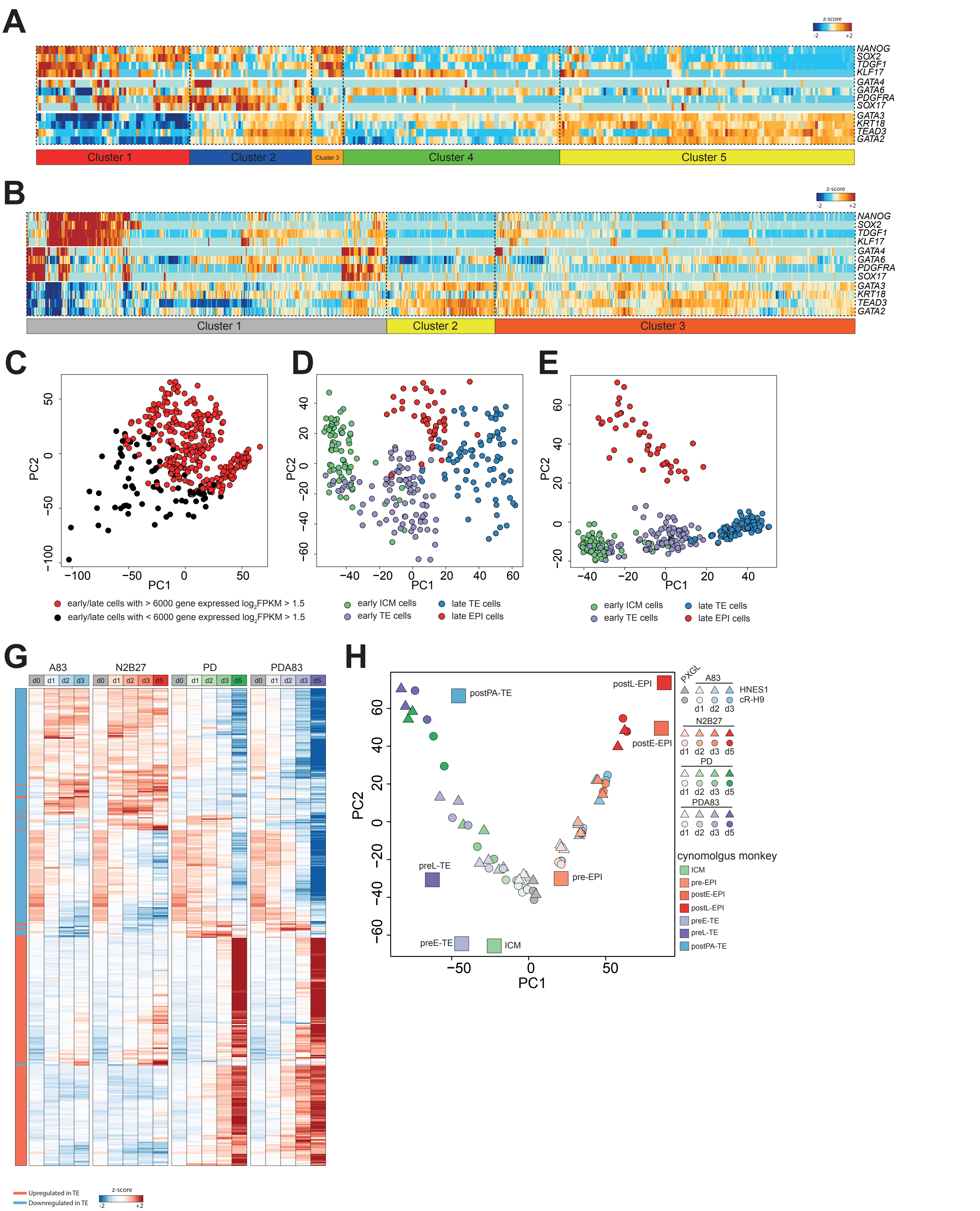
A
z-score
-2
+2
NANOG
SOX2
TDGF1
KLF17
GATA4
GATA6
PDGFRA
SOX17
GATA3
KRT18
TEAD3
GATA2
Cluster 1
Cluster 2
Cluster 3
Cluster 4
Cluster 5
B
Cluster 1
Cluster 2
Cluster 3
C
PC2
PC1
early/late cells with > 6000 gene expressed log2FPKM > 1.5
early/late cells with < 6000 gene expressed log2FPKM > 1.5
D
early ICM cells
early TE cells
late TE cells
late EPI cells
E
G
A83
N2B27
PD
PDA83
d0
d1
d2
d3
d5
Upregulated in TE
Downregulated in TE
H
postPA-TE
postL-EPI
postE-EPI
preL-TE
pre-EPI
preE-TE
ICM
PXGL
HNES1
cR-H9
cynomolgus monkey
ICM
pre-EPI
postE-EPI
postL-EPI
preE-TE
preL-TE
postPA-TE

Figure S4

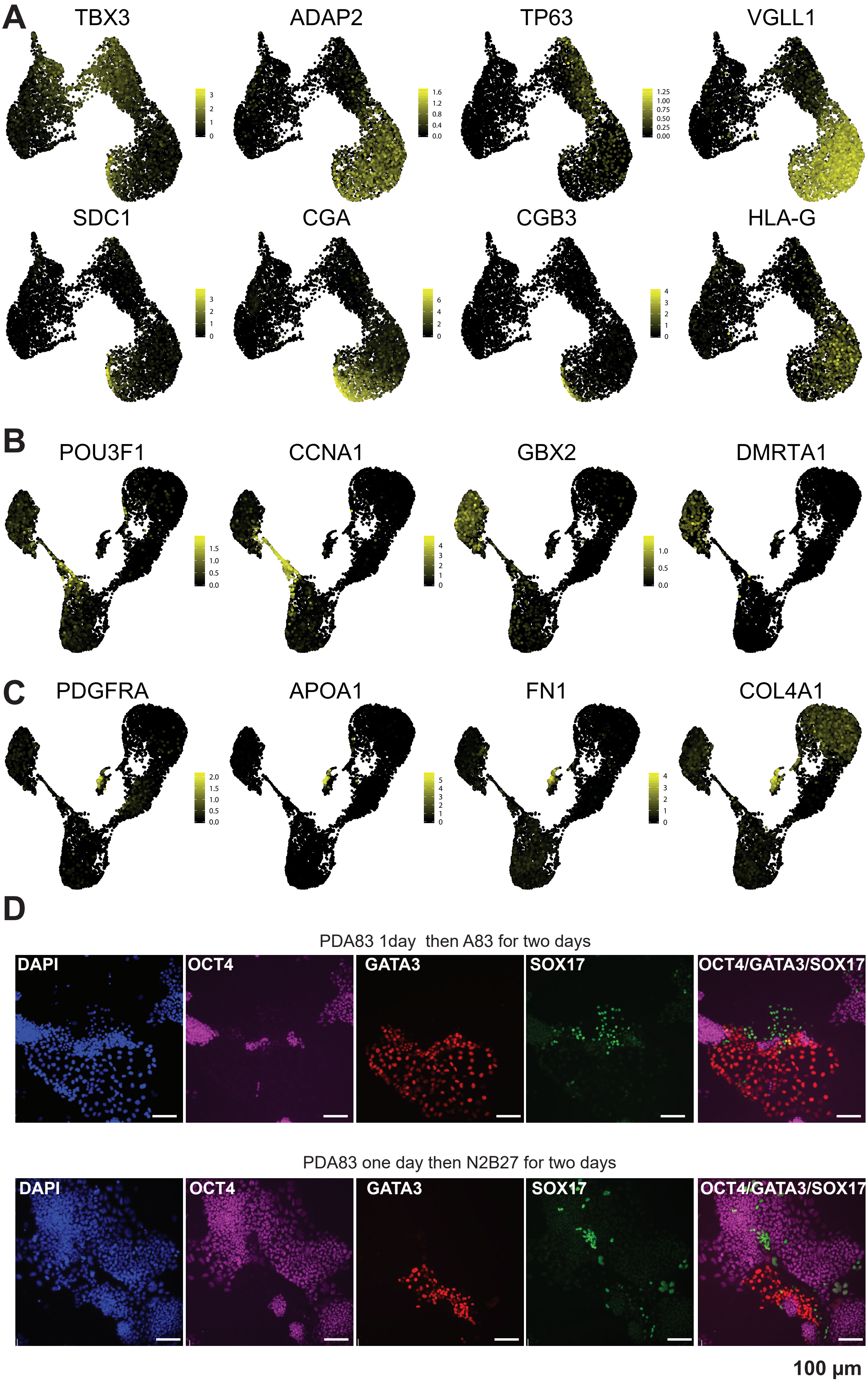
A
TBX3
ADAP2
TP63
VGLL1
SDC1
CGA
CGB3
HLA-G
B
POU3F1
CCNA1
GBX2
DMRTA1
C
PDGFRA
APOA1
FN1
COL4A1
D
PDA83 1day then A83 for two days
DAPI
OCT4
GATA3
SOX17
OCT4/GATA3/SOX17
PDA83 one day then N2B27 for two days
DAPI
OCT4
GATA3
SOX17
OCT4/GATA3/SOX17
100 µm

### Figure S5

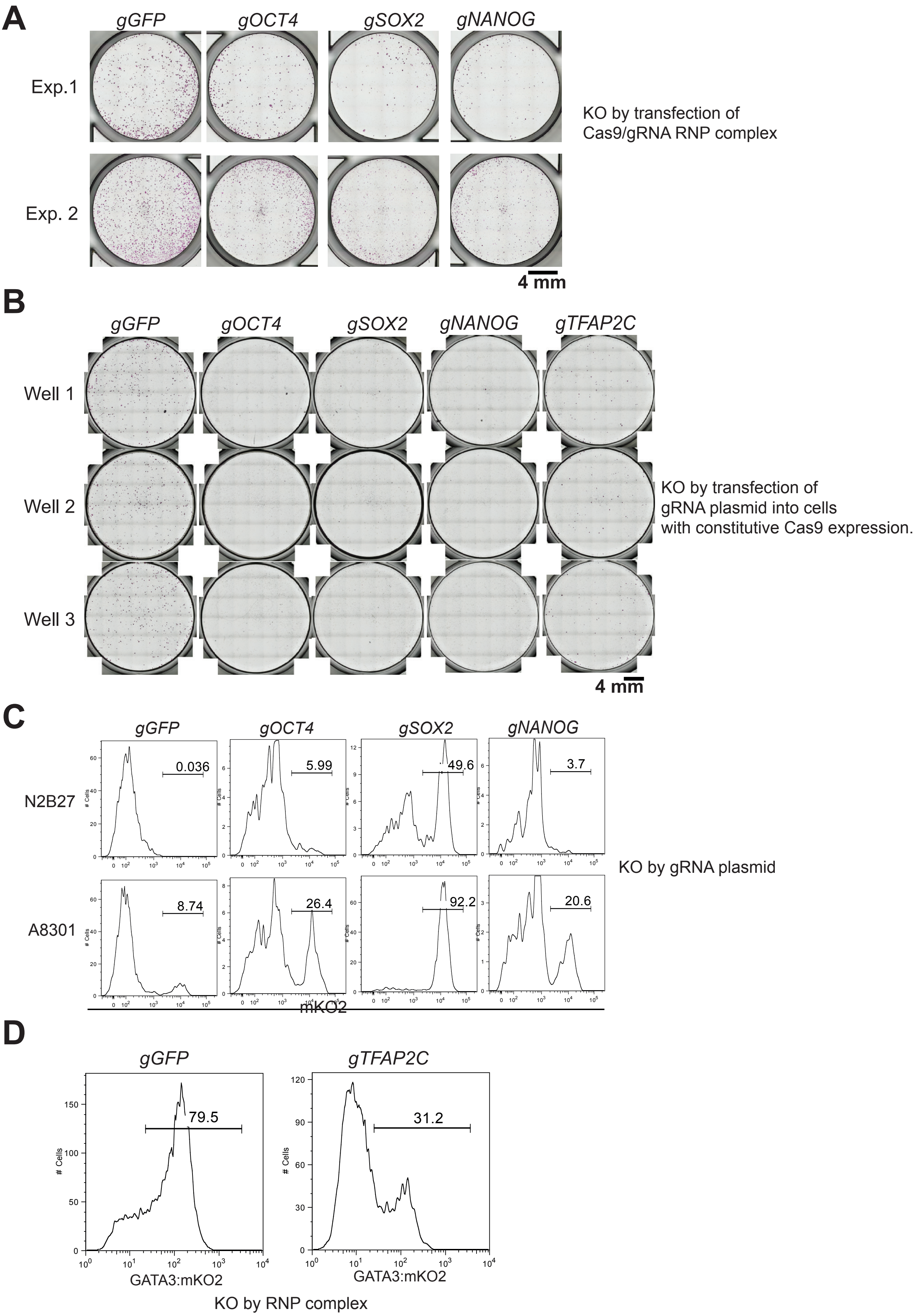
A
gGFP
gOCT4
gSOX2
gNANOG
Exp.1
Exp. 2
KO by transfection of
Cas9/gRNA RNP complex
4 mm
B
gGFP
gOCT4
gSOX2
gNANOG
gTFAP2C
Well 1
Well 2
Well 3
KO by transfection of
gRNA plasmid into cells
with constitutive Cas9 expression.
4 mm
C
gGFP
gOCT4
gSOX2
gNANOG
N2B27
A8301
0.036
5.99
49.6
3.7
8.74
26.4
92.2
20.6
### Cells
mKO2
KO by gRNA plasmid
D
gGFP
gTFAP2C
79.5
31.2
### Cells
GATA3:mKO2
GATA3:mKO2
KO by RNP complex

### Figure S6

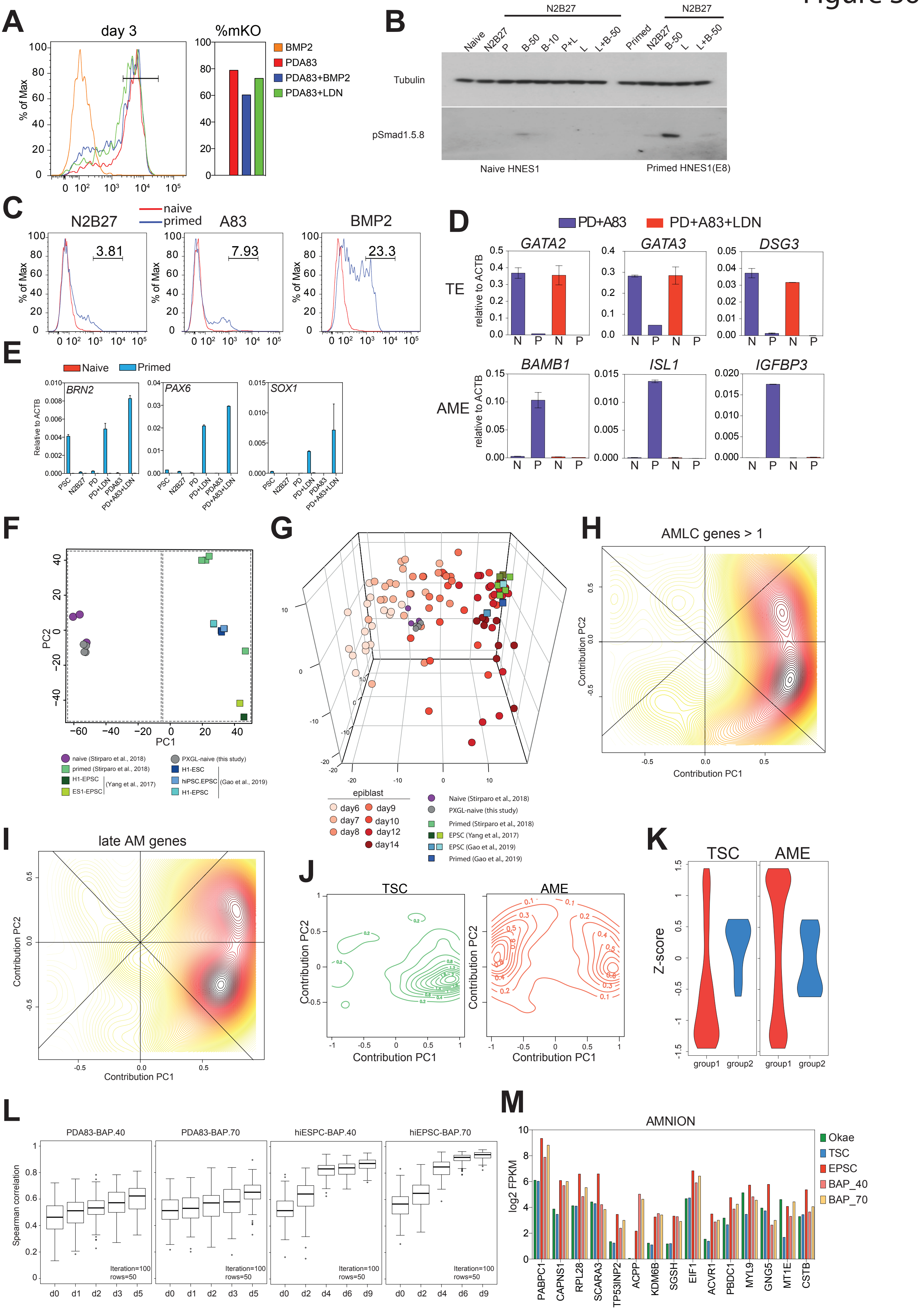
A
day 3
%mKO
% of Max
BMP2
PDA83
PDA83+BMP2
PDA83+LDN
B
N2B27
Naive
Primed
Tubulin
pSmad1.5.8
Naive HNES1
Primed HNES1(E8)
C
N2B27
A83
BMP2
naive
primed
3.81
7.93
23.3
D
PD+A83
PD+A83+LDN
GATA2
GATA3
DSG3
TE
relative to ACTB
BAMB1
ISL1
IGFBP3
AME
E
Naive
Primed
BRN2
PAX6
SOX1
Relative to ACTB
PSC
N2B27
PD
PD+LDN
PDA83
PD+A83+LDN
F
PC1
PC2
naive (Stirparo et al., 2018)
primed (Stirparo et al., 2018)
H1-EPSC
ES1-EPSC
(Yang et al., 2017)
PXGL-naive (this study)
H1-ESC
hiPSC.EPSC
H1-EPSC
(Gao et al., 2019)
G
epiblast
day6
day7
day8
day9
day10
day12
day14
Naive (Stirparo et al., 2018)
PXGL-naive (this study)
Primed (Stirparo et al., 2018)
EPSC (Yang et al., 2017)
EPSC (Gao et al., 2019)
Primed (Gao et al., 2019)
H
AMLC genes > 1
Contribution PC1
Contribution PC2
I
late AM genes
J
TSC
AME
K
TSC
AME
Z-score
group1
group2
L
PDA83-BAP.40
PDA83-BAP.70
hiESPC-BAP.40
hiEPSC-BAP.70
Spearman correlation
Iteration=100
rows=50
d0
d1
d2
d3
d5
d4
d6
d9
M
AMNION
log2 FPKM
Okae
TSC
EPSC
BAP_40
BAP_70
PABPC1
CAPNS1
RPL28
SCARA3
TP53INP2
ACPP
KDM6B
SGSH
EIF1
ACVR1
PBDC1
MYL9
GNG5
MT1E
CSTB

### Figure S7

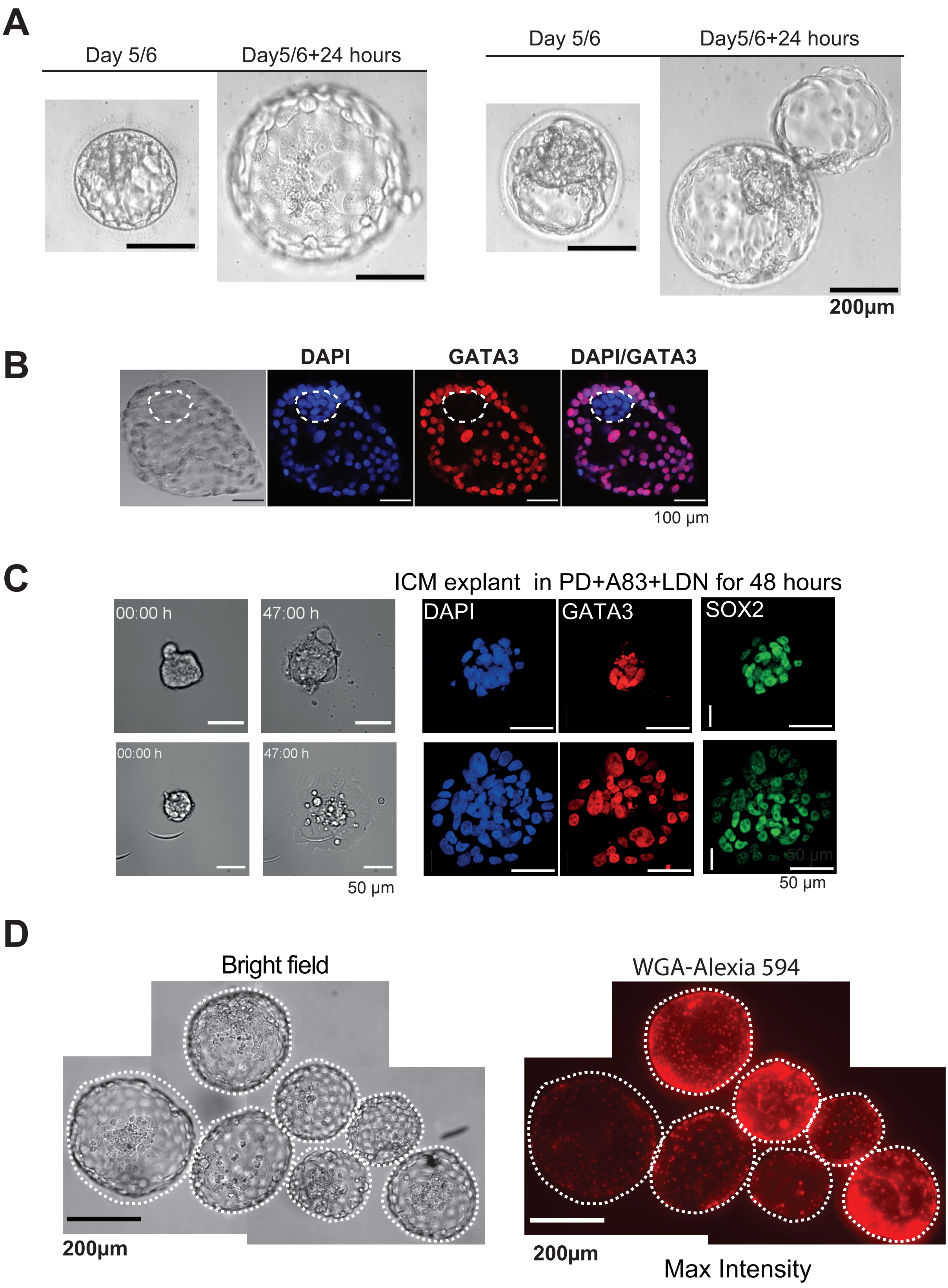
A
Day 5/6
Day5/6+24 hours
Day 5/6
Day5/6+24 hours
200µm
B
DAPI
GATA3
DAPI/GATA3
100 µm
C
ICM explant in PD+A83+LDN for 48 hours
00:00 h
47:00 h
DAPI
GATA3
SOX2
00:00 h
47:00 h
50 µm
50 µm
D
Bright field
WGA-Alexia 594
200µm
200µm
Max Intensity
